# Aperiodic and Hurst EEG exponents across early human brain development: a systematic review

**DOI:** 10.1101/2024.02.02.578622

**Authors:** R. A. Stanyard, D. Mason, C. Ellis, H. Dickson, R. Short, D. Batalle, T. Arichi

## Abstract

In electroencephalographic (EEG) data, power-frequency slope exponents (1/*f*^β^) can provide non-invasive markers of *in vivo* neural activity excitation-inhibition (E:I) balance. E:I balance may be altered in neurodevelopmental conditions; hence, understanding how 1/*f*^β^ evolves across infancy/childhood has implications for developing early assessments/interventions. This systematic review (PROSPERO-ID: CRD42023363294) explored the early maturation (0-26yrs) of resting-state EEG 1/*f* measures (aperiodic [AE], power law [PLE] and Hurst [HE] exponents), including studies containing ≥1 1/*f* measures and ≥10 typically developing participants. Five databases (including Embase and Scopus) were searched during March 2023. Forty-two studies were identified (N_participants_=3478). Risk of bias was assessed using the Quality Assessment with Diverse Studies tool. Narrative synthesis of HE data suggests non-stationary EEG activity occurs throughout development. Age-related trends were complex, with rapid decreases in AEs during infancy and heterogenous changes thereafter. Regionally, AE maxima shifted developmentally, potentially reflecting spatial trends in maturing brain connectivity. This work highlights the importance of further characterising the development of 1/*f* measures to better understand how E:I balance shapes brain and cognitive development.

## Introduction

The maintenance of excitation and inhibition (E:I) balance in the brain is an essential homeostatic mechanism that regulates spontaneous neural activity and facilitates the complex activity patterns thought to underlie efficient information processing (Lendner *et al*., 2020, Weber, Klein and Abeln, 2020, Tran *et al*., 2020) and adaptive behaviour (Rocha *et al*., 2018; Bassi *et al*., 2019). This key feature of brain physiology can be represented by a power law (1/*f*) relationship between spectral frequencies and spectral power in electrophysiological data (Gao, Peterson and Voytek, 2017). Steeper 1/*f* profiles (higher exponents) characterised within specific frequency ranges (Manning *et al*., 2009; Miller *et al*., 2012) suggest higher contributions of inhibitory (*i.e.* increased GABAergic/decreased glutamatergic) signalling whereas flatter (lower) exponents suggest excitation-dominant signalling (E>I) (Gyurkovics *et al*., 2022). This can be non-invasively studied using electroencephalography [EEG] (Waschke, Wöstmann and Obleser, 2017) (Waschke, Wöstmann and Obleser, 2017) which is sensitive to local field potential (LFP) aggregates; and thus changes in power spectral densities (PSDs) will affect estimated 1/*f* exponents. The power spectrum can be further decomposed into both frequency-specific ‘periodic’ oscillations and an ‘aperiodic’ signal (termed *β* or χ) (Voytek *et al*., 2015). In adulthood, *β* significantly declines with age (Voytek, *et al*., 2015; Waschke, Wöstmann and Obleser, 2017) although the physiological origin of this age-related change is unclear. EEG 1/*f* measures also display behavioural and clinical relevance, particularly in conditions thought to relate to shifts in E:I balance, such as those affecting attention and behaviour (Waschke *et al*., 2021; Robertson *et al*., 2019), states of consciousness (Leroy *et al*., 2023), and functional recovery from stroke (Leemburg *et al*., 2018). Prior to future research utilising 1/*f* measures to explore possible atypical brain E:I or as a biomarker in clinical populations, we must first characterise 1/*f* measures across the typically developing (TD) lifespan, from infancy to early adulthood (other studies have begun to chart this for later adulthood, see Finley *et al*., 2022).

Three different methods for deriving the 1/*f*^β^ exist in the human EEG literature: (1) power law exponents (PLEs, He, 2014) estimated from the slope of log-frequency versus log-power distributions and measures accounting for periodic oscillations, including aperiodic exponents (μV^2^ Hz^−1^) [AEs]; (2) fitting of one-over-f [FOOOF] (now specparams) via estimation of an initial slope and iterative estimation of gaussian peaks, which are subsequently subtracted to facilitate slope re-estimation prior to combining into a representative model (Donoghue *et al*., 2020); or (3) from non-integer resampling (Irregular Resampling Auto-Spectral Analysis [IRASA])(Wen and Liu, 2016). Given the challenges of comparing raw exponents acquired when performing different tasks (Gao, Peterson and Voytek, 2017), we focus here only on characterising resting-state 1/*f*^β^ during typical development and maturation. We also explore evidence surrounding the maturation of activity patterns in the temporal domain via the resting Hurst exponent (HE), typically calculated via detrended fluctuation analysis (DFA)(Peng *et al*., 1994, 1995). HE (α) can be converted into PLE for both stationary (α = 0-1) and non-stationary (α = 1-2) cases (Eke *et al*., 2000; Hardstone *et al*., 2012). Given the dynamic nature of the brain’s activity, EEG data generally display persistent patterns of electrical activity (0.50<*HE*<1.00) which are non-stationary (HE>0.50) *i.e.* activity does not revert to a baseline state but is segregated and maintained in contextual functional states. To further synergise the 1/*f* literature here we also convert HE into AE, wherein AE=2*HE-1, thus providing a comprehensive account of early developmental 1/*f*^β^ changes.

This systematic review aims to explore how and when 1/*f* measures change in early human development, and where variability within early lifespan stages exists, thereby offering a more nuanced perspective of sensitive periods of neurodevelopment.

## Methods

### Eligibility Criteria and Selection Process

We included observational or experimental studies containing resting-state (eyes open [EOR] or closed [ECR]) data for ten or more typically developing (TD) human participants with a mean-centred age less than 26.50yrs (*i.e.* bordering into ‘emerging adulthood’, see Hochberg and Konner, 2020). For subjects younger than 2yrs (infants and neonates), data collected during sleep or wake (including when observing videos or toys) were included. We included studies which referred to AE or slope, 1/*f*^β^, HE, fractal dimension (to assess for HEs), PLE/spectral slopes, or AE/PLE estimation models (*e.g.* FOOOF/specparams/IRASA/sprintf/PaWNextra). Abstracts fitting these criteria were assessed as full-texts if an English-language text was available, including abstracts referring to an evoked paradigm or where sample or method details were omitted, so as to capture suitable studies containing resting-state data. Articles focusing on non-human populations (*e.g.* animals, or simulations only), of an unsuitable format (preprints, reviews, theses, case reports, books, conference abstracts, and non-peer-reviewed material) or using measures other than scalp-based EEG (*e.g.* iEEG, sEEG or ECoG, MEG, TMS, tDCS) were excluded. Articles lacking measures of interest in the main/supplementary texts were excluded. In calculating the HE, the underlying scaling exponent (α) only deviates from 0.50 for short window sizes (see Hardstone *et al*., 2012), hence the scaling range should be reported. Furthermore, we exclude papers not reporting or responding to requests for two or more key details (scaling range, epoch length, window size).

### Search Strategy and Information Sources

The systematic review was completed according to the PRISMA guidelines (Page *et al*., 2021) and pre-registered with the international Prospective Register of Systematic Reviews (PROSPERO Registration number: CRD42023363294). Relevant literature referred to the development/maturation of the 1/*f*^β^ signal: 1/*f*, aperiodic exponent/slope and/or the HE (Hurst exponent*/slope or fractal, primarily measured via DFA or detrend* fluctuation analysis) across the early human lifespan (birth, newborn, neonat*, infan*, toddler*, child*, adolescent, teenager, young adult*, develop*, maturation*) as measured using EEG (EEG or electroencephal*). Searches were performed across the following databases (with appropriate MESH headings and adjacency terms where permissible): Ovid-Embase, Ovid-PsycInfo, Ovid-Medline, Scopus and Web of Science, during March 2023. For an example search strategy, see **Supplementary Material I**. Backwards searching of included studies was also performed.

### Selection Process

Records were stored and de-duplicated in Endnote before being transferred to Rayaan for secondary deduplication and subsequent screening. Titles and abstracts were screened by author RAS, with a subset (20%) reviewed independently by co-author DM and re-reviewed in cases of disagreement until a consensus was reached.

### Data Collection and Data Items

Article full texts were then screened by RAS and data pertaining to sample characteristics (age mean and SD, sample size, gender split) and 1/*f* data (AE/PLE/HE) were extracted from tables or figures of the main and/or supplementary texts, an associated repository or by contacting the authors directly.

### Study Risk of Bias Assessment

Risk of bias was assessed independently by co-authors RAS and CE using the Quality Assessment for Diverse Studies (QuADS) tool (Harrison *et al*., 2021), with the omission of item 12 (stakeholder involvement) due to a lack of relevance to the TD population. Rater scores (91.07% agreement) were compared to ensure differences of less than 2 points (0.01%, 6/504 cases). Differing cases were discussed, agreed and calibrated. For the assessment criteria and risk of bias results, see **Supplementary Materials II and III** respectively.

### Synthesis Methods

Few studies reported age correlations or other effect sizes (N=8) and given the ambiguity of raw AE effect size interpretation (Gao, Peterson and Voytek, 2017) and the absence of a comparison state uniform to all studies, a meta-analysis was not performed. Rather, we qualitatively synthesised findings across lifespan stages: infancy (0.01-2.00yrs), toddlerhood (2.00-3.00yrs), childhood (3.00-12.99yrs), adolescence (13.00-19.99yrs), young adulthood (20.00-26.00yrs), spatial scale (global, regional, channel-wise), method (HE, PLE/AE) and condition (ECR/EOR).

## Results

Our database searches yielded 1,596 records. After de-duplication, we screened 1,112 titles and abstracts. Nine full-texts sought for retrieval were unavailable, resulting in 138 retrieved full-texts, of which 37 were included (see **Figure 1**). We identified a further five studies after searching for citations of the included studies as well as their reference lists. The characteristics of the included studies are shown in **Table 1**.

**Figure 1.**
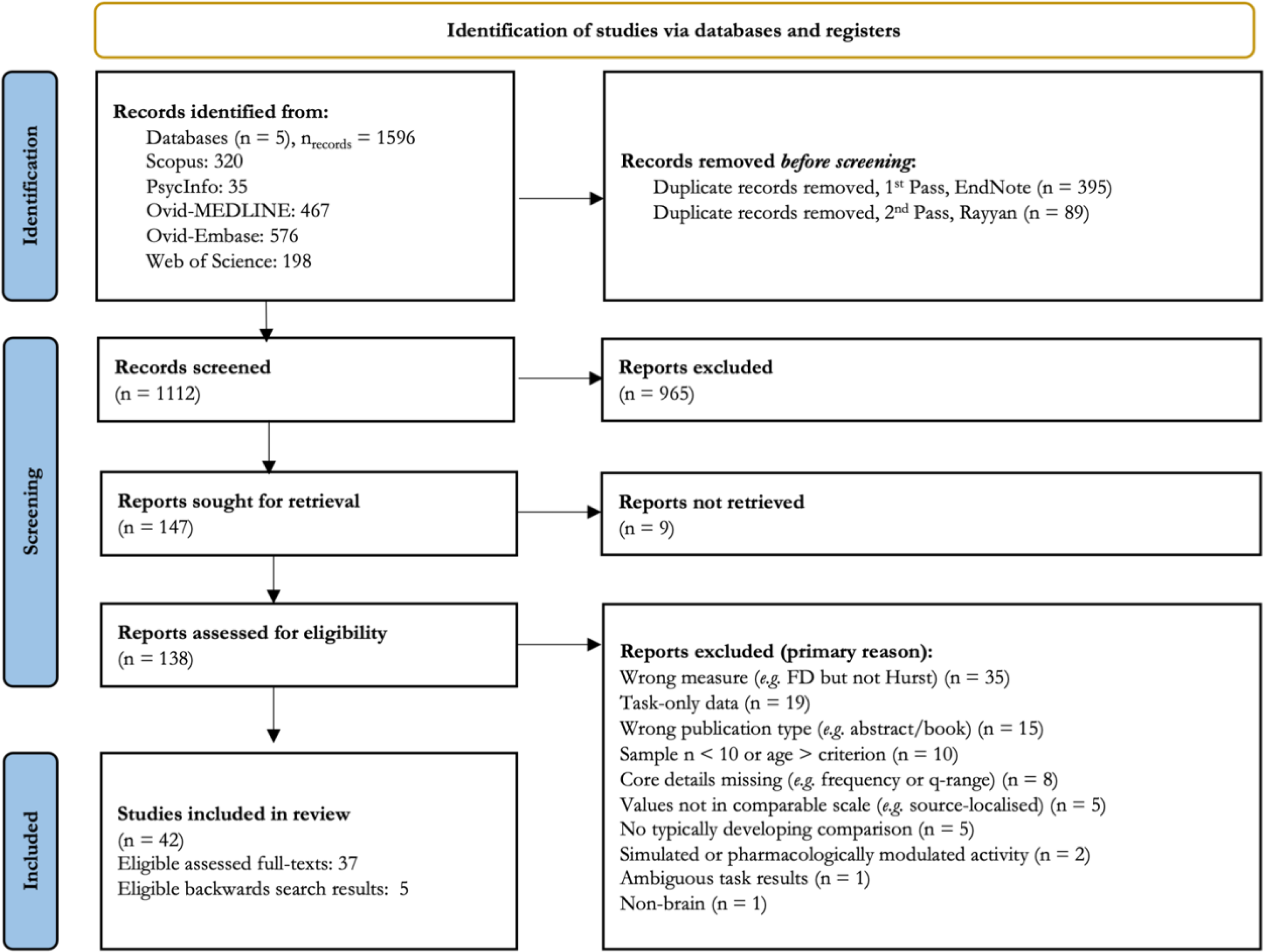
PRISMA flowchart for record screening. Backwards searching utilised based on citation title relevance of included texts to ensure sufficient article capture (N=5 relevant reports, see ‘Included’).

**Table 1.** Studies included in the review. Infancy (0.01-2.00yrs), Toddlerhood (2.00-3.00yrs), Childhood (3.00-12.99yrs), Adolescence (13.00-19.99yrs), Young adulthood [YA] (20.00-26.00yrs). ‘^est^’ in the ‘Scale’ column denotes values are estimated from a plot. Measures include eyes open (EOR) and closed (EOR) rest, alongside other specified states. Measures from sub-samples in the ‘Original Measure’ column are referred to by ‘S’ whilst observed timepoints are denoted by ‘t’. Sample split by sex is given in ‘N (M, F)’, wherein unknown values are indicated by ‘?’. Data from supplementary sources (tables, figures) are denoted as ‘Supp.’ in the ‘Source’ column, with open-access data from the open science framework (OSF) marked and ‘Auth Corr.’ denoting author correspondence was required for additional information/data was absent from the published material. Sections and figures (‘Fig’) are marked where relevant. Studies with overlapping data are marked with the same superscript character (^a,b^ respectively). In column ‘F’, ‘Y’ entries denote backward-search results. For technical details of measures, see **Supplement IV**.

| # | Study | Lifespan Stage (age, yrs) | Measure | Scale(s) | Original Measure | HE to AE | Measure | N (M, F) | Source | F |
| --- | --- | --- | --- | --- | --- | --- | --- | --- | --- | --- |
| 1 | Schaworonkow & Voytek (2021) | Infancy (0.10-0.56) | 1/f (FOOOF) | Channelwise | S1: 1.74-3.22 (N = 20)<br>S2: 1.74-2.95 (N = 20)<br>S3: 1.79-2.25 (N = 20)<br>S4: 1.94-2.98 (N = 5)<br>S5: 1.46-2.76 (N = 3)<br>S6: 1.88-2.63 (N = 2) |  | Baseline-wakeful reaching | 22(10,12) | Methods, Auth Corr., GitHub |  |
| 2 | Karalunas (2022) | Infancy (0.12±0.01)<br>Adolescent (14.10±1.30) | 1/f (FOOOF) | Global,<br>Channelwise | Infant EOR (PEACH cohort): 2.21±0.28<br>Adol EOR (1.85±0.28): ECR (1.98±0.26),<br>EOR-ECR avg (1.91±0.28)<br>Infant: 2.48±0.24 (Cz, EOR)<br>Adol: 2.28±0.19 (Cz, EOR),<br>2.33±0.27 (Pz, ECR),<br>2.29±0.19(EOR-ECR average) |  | EOR, ECR<br>EOR-ECRavg | 69 (36,33)<br>152(85,67) | Auth Corr. |  |
| 3 | Fransson (2013) | Infancy (0.81, 0.75-0.85) | 1/f (PLE) | Global,<br>Regional,<br>Channelwise | 2.07±0.22 |  | Natural Active/<br>Quiet Sleep | 15(9,12) | Fig 4 | Y |
| 4 | Carter-Leno (2022) | Infancy (0.90±0.05) | 1/f (FOOOF) | Global,<br>Regional,<br>Channelwise | 1.50±0.13 (non-social), 1.52±0.16<br>Fz: social (1.53±0.16), non-social (1.51±0.13)<br>Cz: social (1.51±0.15), non-social (1.49±0.13)<br>Pz: social (1.51±0.16), non-social (1.49±0.13) |  | ~EOR (social and non-social videos) | 24(13,11) | Table 1, Fig 4, Auth Corr. |  |
| 5 | Roche (2019) | Infancy (1.92-10.25) | 1/f (PLE) | Global<sup>est</sup>,<br>Regional | ~0.58 |  | ~EOR (movie) | 37(0,37) | Methods, Results |  |
| 6 | Smith (2021) | Infancy (med. 0.63, 0.43-0.82) | HE | Global | Delta[1-3Hz]: ~0.80 (A), 0.68(S)<br>Theta[4-7Hz]: ~0.74(A), 0.68(S)<br>Alpha[8-12Hz]: ~0.69(A), 0.68(S)<br>Beta[13-30Hz]: ~0.88(A), 0.72(S) |  | Awake, Sleep | 20(12,8) | Section 3.1, Fig 6, Auth Corr. |  |
| 7 | Smith (2017) | Infancy (med. 0.58, 0.48-0.94) | HE | Global<sup>a</sup> | Delta[1-3Hz]: ~0.78<br>Theta[4-7Hz]: ~0.70<br>Alpha[8-12Hz]: ~0.66<br>Beta[13-30Hz]: ~0.94 |  | Awake<br>~(EOR) | 21(?,?) | Fig 5, Auth Corr. |  |
| 8 | Cellier (2021) | Toddler (N=5),<br>Child (N=81),<br>Adolescent (N=22),<br>Young Adult (N=8)<br>(2.95-24) | 1/f<br>(FOOOF) | Regional<br>(Parietal-midline<br>[P], Frontal-<br>midline [F]) | Toddler: [P] 1.45±0.23, [F] 1.32±0.54<br>Children: [P] 1.23±0.25, [F] 1.34±0.22<br>Adolescents: [P] 1.24±0.18, [F] 1.13±0.24<br>Young Adults: [P] 1.14±0.12, [F] 1.11±0.09 |  | EOR | 116 (33,24,59<br>unlabelled) | Fig 2,<br>Sections 2.2,<br>3.1,<br>Auth Corr.,<br>OSF |  |
| 9 | Houtman (2021) | Toddler (2.92 [N=8], 3.92<br>[N=13]),<br>Child<br>(7-16[N=29]) | 1/f<br>(FOOOF),<br>HE | Global<br>Channelwise | Infant-toddler (I) & child-adol (C): HE, 11-18Hz: I: ~0.655, C: ~0.656<br>Infant-toddler (I) & child-adol (C): AE,<br>~1.11-1.60<br>(Hurst, 11-18Hz): I: ~0.63-.70, C: ~0.64 -<br>0.74, 0.66±0.02 |  | EOR | 50 (28,22):<br>Inf-Todd:<br>21 (14,7)<br>Child-Adol:<br>29 (14,15)<sup>a</sup> | Fig 3, 5<br>Supp. Fig 5 |  |
| 10 | Wilkinson & Nelson<br>(2021) | Child<br>(3.98±1.09, 2.67-6.67) | 1/f<br>(FOOOF) | Global<br>Regional | 1.19±0.12<br>Frontal: 1.26±0.13<br>Central: 1.33±0.14<br>Temporal: 1.11±0.15<br>Posterior: 1.07±0.32 |  | EOR | 12(12,0) | Methods,<br>Results,<br>Auth Corr. |  |
| 11 | Robertson (2019) | Child<br>(5.65±1.23) | 1/f<br>(FOOOF) | Global<br>Channelwise | 1.51±0.32 |  | EOR | 50(36,14) | Table 1,<br>Fig 2A, B |  |
| 12 | McSweeney (2023) | Child<br>(6.92±2.21) | 1/f<br>(FOOOF) | Global | EOR: 1.53±0.31<br>ECR: 1.77±0.28 |  | EOR, ECR | 502(230,272) | Section 3.2,<br>Auth Corr. |  |
| 13 | Arnett (2022a)<br>...Stein | Child<br>(8.83±1.23) | 1/f<br>(FOOOF) | Global | 1.77±0.15 (Median: 1.76) |  | EOR | 29(19,10)<sup>b</sup> | Methods,<br>Auth Corr. |  |
| 14 | Arnett (2022b)<br>... Levin | Child<br>(8.83±1.23) | 1/f<br>(FOOOF) | Global | 1.77±0.15 (Median: 1.76, range: 0.22-2.30) |  | EOR | 29(19,10)<sup>b</sup> | Methods,<br>Auth Corr. |  |
| 15 | Peisch & Arnett<br>(2022) | Child<br>(9.40±1.36) | 1/f<br>(FOOOF) | Global<br>Regional | 1.78±0.14<br>Anterior Frontal (AF): 1.79±0.14<br>Frontal (FR): 1.79±0.13<br>Central (CE): 1.75±0.15<br>Parietal (PR): 1.81±0.16<br>Occipital (OC): 1.77±0.22 |  | EOR | 29(19,10)<sup>b</sup> | Methods,<br>Auth Corr. |  |
| 16 | Hill (2022) | Child<br>(9.41±1.95) | 1/f<br>(FOOOF) | Global<br>Regional<br>(anterior [A],<br>central [C],<br>posterior [P]) | EOR: 1.65±0.18<br>ECR: 1.81±0.16<br>EOR: A (1.64±0.19), C (1.69±0.19)<br>P (1.68±0.20)<br>ECR: A (1.81±0.17), C (1.85±0.16),<br>P (1.84±0.18) |  | EOR, ECR | 139 (72, 67) | Fig 2,<br>Auth Corr. |  |
| 17 | Tröndle (2022) | Child (N=153),<br>Adolescent (N=34),<br>Young Adult (N=3)<br>(10.07±3.39, 5.02-21.67) | 1/f<br>(FOOOF) | Regional<br>(Parieto-<br>occipital) | 1.89±0.36 (0.68-2.77)<br>Child: 1.98±0.30<br>Adolescent: 1.58±0.37<br>Young adult: 1.12±0.04 |  | ECR | 190<br>(104,86) | Methods,<br>Auth Corr.,<br>Fig 3, App. 4,<br>Supp. 2, 3 |  |
| 18 | Kwok (2019) | Child<br>4yrs (N=8),<br>5yrs (N=14),<br>6yrs (N=11),<br>(5.60±?.??) | HE | Global,<br>Channelwise<sup>est</sup> | Median: ~0.09 (EOR) ~0.06(ECR)<br>Posterior electrodes: ~0.09 EOR, ECR |  | EOR, ECR | 33(?,?) | Fig 6A-C |  |
| 19 | Smit (2011) | Child (5.27±0.19,<br>6.79±0.19),<br>Adolescent (16.06±0.55<br>17.57±0.55), | HE | Channelwise (12<br>channels) | Child Theta (P3 maxima): 0.77±0.09 (5yrs),<br>0.76±0.07 (7yrs)<br>Child Alpha (O2 maxima): 0.70±0.09 (5yrs),<br>0.71±0.08 (7yrs) |  | ECR | 5yrs 366<br>7yrs 378<br>16yrs 426 | Auth Corr.<br>Methods,<br>Fig 3,<br>Table 2 |  |
|  |  | Young Adult (26.18±4.15)<br>(5-50) |  |  | Child Beta: 0.64±0.09 (5 yrs, Fp2),<br>0.62±0.08 (7yrs, F8)<br>Adol Theta (Fp1 maxima): 0.72±0.06<br>(16yrs), 0.72±0.06 (18yrs)<br>Adol Alpha (O1 maxima): 0.72±0.10 (16yrs),<br>0.73±0.12 (18yrs)<br>Adol Beta: 0.64±0.09 (16yrs), 0.66±0.11<br>(18yrs)<br>YA Theta (F3 maxima): 0.73±0.07 (25yrs)<br>YA Alpha (P4 maxima): 0.75±0.09 (25yrs)<br>YA Beta (O1 maxima): 0.67±0.10 (25yrs) |  |  | 18yrs 387<br>25yrs 396 |  |  |
| 20 | Bruining (2020) | Child<br>(10.30±1.54) | HE | Global | 0.66±0.04 |  | ECR | 29 (14,15)<sup>a</sup> | Supp. Table 1 | Y |
| 21 | McSweeney (2021) | Adolescent<br>(12-17) | 1/f<br>(FOOOF) | Global (1-45Hz) | t1 (all subjects): EOR (1.21±0.30)<br>t1 (subjects with t1 & t2): EOR (1.21±0.30)<br>t1 (all subjects): ECR (1.33±0.27)<br>t1 (subjects with t1 & t2): ECR (1.21±0.30)<br>t2 (all subjects): EOR (1.10±0.26)<br>t2 (subjects with t1 & t2): EOR (1.11±0.27)<br>t2 (all subjects): ECR (1.16±0.25)<br>t2 (subjects with t1 & t2): ECR (1.17±0.26) |  | EOR, ECR | 186<br>(85,101)<br>95 @t<sub>1</sub>, t<sub>2</sub> | Fig 1B,<br>Results |  |
| 22 | Ostlund (2021) | Adolescent<br>(13.97±1.28) | 1/f<br>(FOOOF) | Global (2-50Hz) | (EOR+ECR/2): 1.80±0.28 (0.92-2.57)<br>EOR: 1.72±0.31 (0.88-2.49)<br>ECR: 1.88±0.28 (0.90-2.65) |  | EOR, ECR | 97(53,43) | Table 1 |  |
| 23 | Linkenkaer-Hansen<br>(2007) | Adolescent<br>(16.50-19.50) | HE | Channelwise<br>(Alpha, Beta) | Alpha: (0.70-0.74±0.08-0.11)<br>Beta: (0.61-0.66±0.07-0.09) |  | ECR | 390<br>(196,194) | Table 1 |  |
| 24 | Gao, F. (2017) | Adolescent<br>(18.30±2.80) | HE | Channelwise<br>(Delta-Gamma)<sup>est</sup> | Alpha: ~0.80<br>Beta: ~0.70 | 0.60<br>0.40 | ECR | 15(15,0) | Fig 2 |  |
| 25 | Donoghue (2020) | Young Adult<br>(19.56±1.90) | 1/f<br>(FOOOF) | Channelwise<br>(Cz) | 1.43±0.25 |  | EOR | 16(8,8) | Auth Corr.,<br>Results |  |
| 26 | Linkenkaer-Hansen<br>(2001) | Young Adult<br>(20-30) | 1/f (PLE)<br>HE | Global (Alpha<br>[8-13Hz])<br>4-channel avg | PLE ECR: 0.36±0.17<br>PLE EOR: 0.51±0.12<br>HE ECR: 0.68±0.07<br>HE EOR: 0.70±0.04 |  | EOR, ECR | 10(9,1) | Results | Y |
| 27 | Muthukumaraswamy<br>& Liley (2018) | Young Adult<br>(23.00±??) | 1/f<br>(IRASA) | Global,<br>Channelwise | $\beta_{lf}$ 1.36(1.12-1.72)<br>$\beta_{hf}$ 1.48(1.18-1.81)<br>$\beta_{lf}$ frontal maxima: 1.72<br>$\beta_{hf}$ central maxima: 1.81 | | ECR | 17(17,0) | Methods,<br>Supp. Fig 7 | |
| 28 | Pathania (2021) | Young Adult<br>(20.88±2.24) | 1/f (PLE)<br>1/f<br>(FOOOF) | Global<br>(FOOOF),<br>Regional<br>(FOOOF), | 1.36±0.26<br>F(1.18±0.34), C(1.40±0.28),<br>P(1.46±0.28), O(1.41±0.29) |  | EOR | 59(19,40) | Auth Corr. |  |
| 29 | Barry and de Blasio<br>(2021) | Young Adult<br>(21.20±3.80) | 1/f PN<br>Slope<br>(PaWNextra) | Global<br>Channelwise (30<br>channels) | EOR (session 1, 2 average): 1.07±0.33<br>ECR: 1.22±0.38<br>EOR: 0.41-1.50 (Fp1, Cz)<br>ECR: 0.38-1.22 (Fp1, C4) |  | EOR, ECR | 20(3,17) | Auth Corr. |  |
| 30 | Merkin (2023) | Young Adult<br>(22.20±3.90,18–35) | 1/f<br>(FOOOF) | Global<sup>est</sup><br>Regional | ~1-2.1<br>~ range 1.3-1.6 YA |  | ECR | 85(37,48) | Sections 2.1,<br>3.1, Supp. S5 |  |
| 31 | Ke (2022) | Young Adult<br>(22.29±2.28) | 1/f<br>(FOOOF) | Global,<br>Regional<br>(Frontal,<br>Central, Parietal,<br>Occipital) | Global (1.84±0.34)<br>Frontal (1.99±0.35)<br>Central (1.84±0.34)<br>Parietal (1.76±0.37)<br>Occipital (1.67±0.52) |  | EOR | 90(44,46) | Table 1,<br>Auth Corr. |  |
| 32 | Smit (2013) | Young Adult<br>(22.40, 21–25) | 1/f (PLE)<br>HE | Channelwise<br>(Alpha [9-13Hz])<br>CP3 | Maxima (both): central midline, scalp ranges<br>Hurst (0.70-0.80), 1/f (0.20-0.40)<br>PLE = 0.43<br>HE = 0.66 (Range: 0.66-1.04) |  | EOR | 39(11,28) | Fig 1B/C,<br>Auth Corr. | Y |
| 33 | Zsido (2022) | Young Adult<br>(22.48±3.79) | 1/f<br>(FOOOF) | Global<sup>est</sup> | ~1.40 |  | ECR | 31(?,?) | Methods |  |
| 34 | Immink (2021) | Young Adult<br>(22.67±3.85) | 1/f<br>(IRASA) | Global | 2.06±0.13 (range: 1.82-2.48) |  | ECR | 45(22,23) | Section 3.1,<br>Auth. Corr. |  |
| 35 | Pathania (2022) | Young Adult<br>(23.29±3.47) | 1/f<br>(FOOOF) | Global (2-25Hz),<br>Regional | 1.17±0.23<br>F(1.20±0.25), C(1.22±0.27), P(1.09±0.28),<br>O(0.96±0.28) |  | ECR | 21(11,10) | Section 4.1,<br>Figure 2B,<br>Auth Corr. |  |
| 36 | Cross (2022) | Young Adult<br>(25.00±7.13) | 1/f<br>(FOOOF,<br>IRASA) | Global | IRASA ECR: 1.11±0.30<br>IRASA EOR: 1.08±0.31 |  | EOR, ECR | 35 (18,17) | Auth Corr. |  |
| 37 | Nakao (2019) | Young Adult<br>(19.57±??<br>18-21) | HE | Alpha [8-13Hz] | FCz: 0.75±0.12<br>Min (T7, 0.74±0.12)<br>Max (O1, 0.80±0.13) | 0.50<br>(0.48-0.60) | ECR | 23(11,12) | Fig 5,<br>Section 3.3,<br>Auth Corr. |  |
| 38 | Natarajan (2004) | Young Adult<br>(20.00±3.00) | HE | Global (1-50Hz) | 0.29±0.06 | -0.42 | ECR | 30(15,15) | Table 1 |  |
| 39 | Liu, S (2022) | Young Adult<br>(20-30) | HE | Channelwise<sup>est</sup><br>(Broadband)<br>[0.5-120Hz] | EOR<br>~0.80-0.82 | EOR<br>0.60-0.64 | ECR, EOR | 26(?,?) | Fig 5 |  |
| 40 | Sleimen-Malkoun<br>(2015) | Young Adult<br>(22.70±1.60, 18.80-25.10) | HE | Global (0.5-100Hz) | 1.69<br>Higher for posterior vs midline | 2.38 | ECR | 31(17,14) | Fig 4 | Y |
| 41 | Irmischer (2018) | Young Adult<br>(25.00±6.20) | HE | Global (Delta<br>[1-4Hz], Theta<br>[4-8Hz], Alpha<br>[8-13Hz], Beta<br>[13-45Hz]) | ECR (N = 57)<br>Theta: 0.66±0.01<br>Alpha: 0.71±0.01<br>Beta: 0.66 ± 0.01<br>EOR (N = 23)<br>Theta: 0.69±0.02<br>Alpha: 0.75±0.02<br>Beta: 0.70 ± 0.01 | ECR<br>0.32<br>0.42<br>0.32<br>EOR<br>0.38<br>0.50<br>0.40 | EOR, ECR | 57(22,35) | Results |  |
| 42 | Bornas (2013) | Young Adult<br>(24.61±7.03) | HE | Regional (theta<br>[3-7Hz], alpha<br>[8-13Hz],<br>broadband [1-40Hz]): Central<br>[C], Parietal [P],<br>Occipital [O]) | Theta<br>C(0.75±0.07), P(0.76±0.07), O(0.74±0.07)<br>Alpha<br>C(0.76±0.07), P(0.80±0.08), O(0.85±0.10)<br>Broadband<br>C(0.85±0.07), P(0.86±0.06), O(0.88±0.06) | Theta<br>0.50,0.52<br>0.48<br>Alpha<br>0.52, 0.60<br>0.70<br>Broadband<br>0.70, 0.72<br>0.76 | EOR, ECR<br>average | 56(20,36) | Table 1 |  |

### Risk of Bias

Across the 12 QuADS items examined, the performance of included studies was generally strong across all items with average scores exceeding 2 (scale 0-3, **Supplementary Material III**). Studies generally showed the weakest performance in terms of providing recruitment data, discussing study strengths and limitations and providing clearly defined research aims/hypotheses.

### Narrative Synthesis

Of the 42 included articles (N=3,478 aggregated observations; 99 HEs+AEs/PLEs, 1097HEs, 2282AEs/PLEs), seven included infants (5 AE/PLE, 2 HE), two included samples containing toddler cohorts (1 AE, 1 AE & HE), thirteen included children (9 AE, 3HE, 1 AE & HE), eight included adolescents (5 AE, 3 HE), and twenty-one included young adults (12 AE, 7 HE and 2 HE and PLE). Most studies analysed data in either EOR or ECR conditions, though two studies used EOR-ECR averages to increase the signal-noise ratio (SNR) (no statistical ECR-EOR differences were reported). The majority of 1/*f*^β^ studies used the FOOOF package (22/42), and thus, for brevity, studies should be assumed to use FOOOF unless otherwise stated. Results are discussed as measured (*i.e.* HE as HE, not AE), with later discussion on the utility of value conversion (see also **Table 1**).

Overall, the method employed to measure 1/*f*^β^ only has a marked impact when comparing converted HE with measurements of AE/PLE, whilst comparisons of direct measures (*i.e.* measures not converted from HEs) show no difference between calculation methods (**Figure 2A**). Focusing on direct AE measures, the global AE decreases from infancy to toddlerhood and remains within more confined AE ranges thereafter (**Figure 2B**). However, the interpretation of this trajectory hinges on an accurate characterisation of AEs during infancy (via sufficiently powered studies), whereas currently, few studies exist. Further, there does not appear to be a difference between global versus regional AEs across the lifespan (**Figure 3A**), evident also on a regional scale (**Supplement V**). Both ECR and EOR AEs display broad variability (**Figure 2B**), particularly in young adulthood (YA), irrespective of study size. Following infancy, regional and global age-related changes generally overlap, with the highest (global) between-study variability observed in YA. These data suggest no differences between AE estimation method, resting-state paradigm, or the level of scale measurement (for most stages). Given the comparability of EOR and ECR, we plot results only for EOR where study data for both conditions is available.

**Figure 2.**
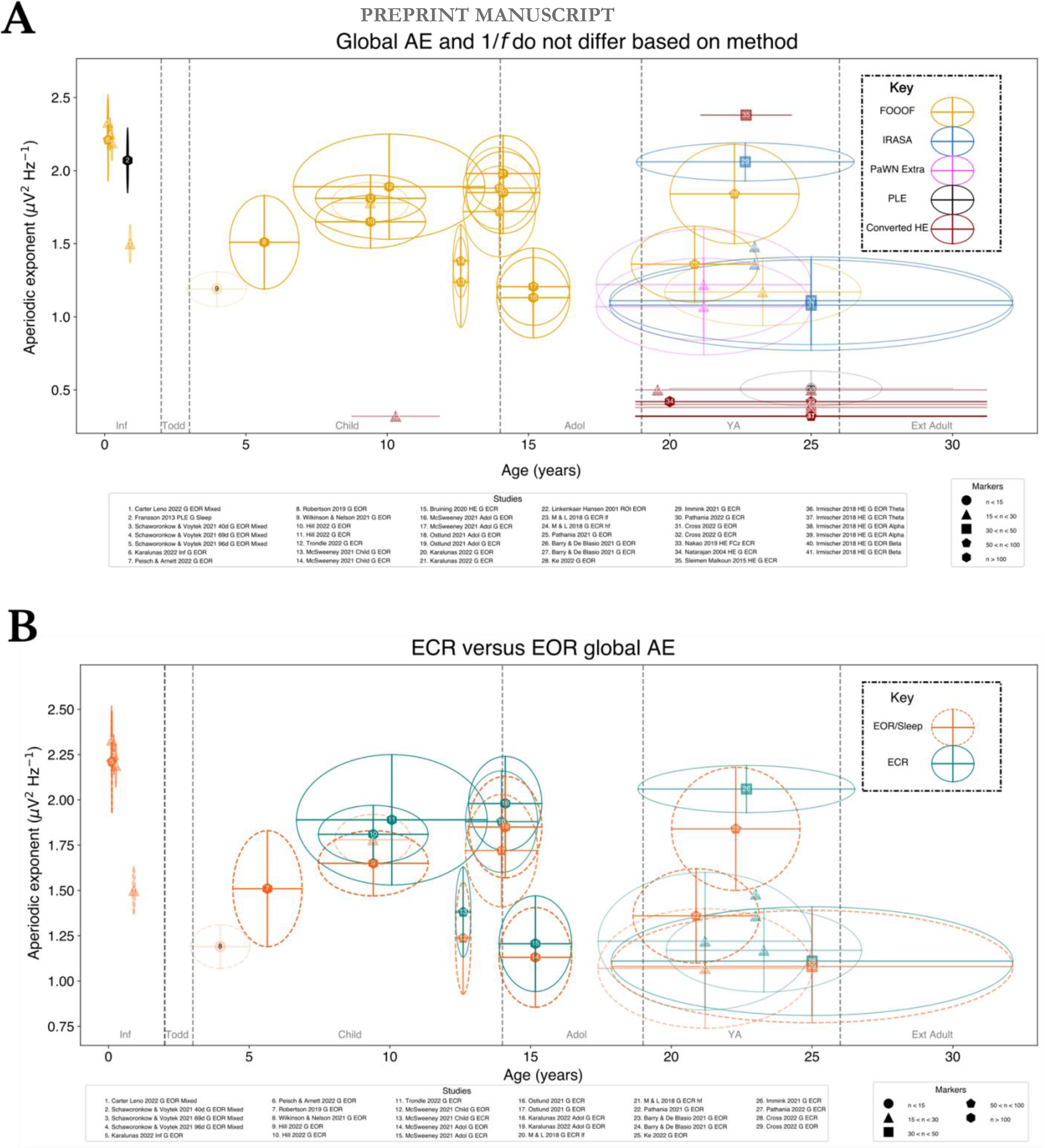
Consistency of the (A) global aperiodic exponent (μV2 Hz−1) across methods and (B) resting-state method for each lifespan stage. Studies are denoted beneath each plot, both figures include eyes open (EOR) and closed (ECR) rest; larger samples are encoded with higher alpha in each plot; see the marker legend for corresponding glyphs. Inf: Infancy, Todd: Toddlerhood, Child: Childhood, Adol: Adolescence, YA: Young adulthood, Ext Ad: Extended adulthood. Horizontal whiskers denote study age standard deviation (SD) whilst vertical whiskers denote 1/*f*^β^ SD. Study numbers (white, black) only differ to enhance readability. ‘lf’ and ‘hf’ denote low and high frequency slope estimation ranges. For visibility, only global AEs are shown, whilst converted HE may include regional measures as the only recording sites available.

**Figure 3.**
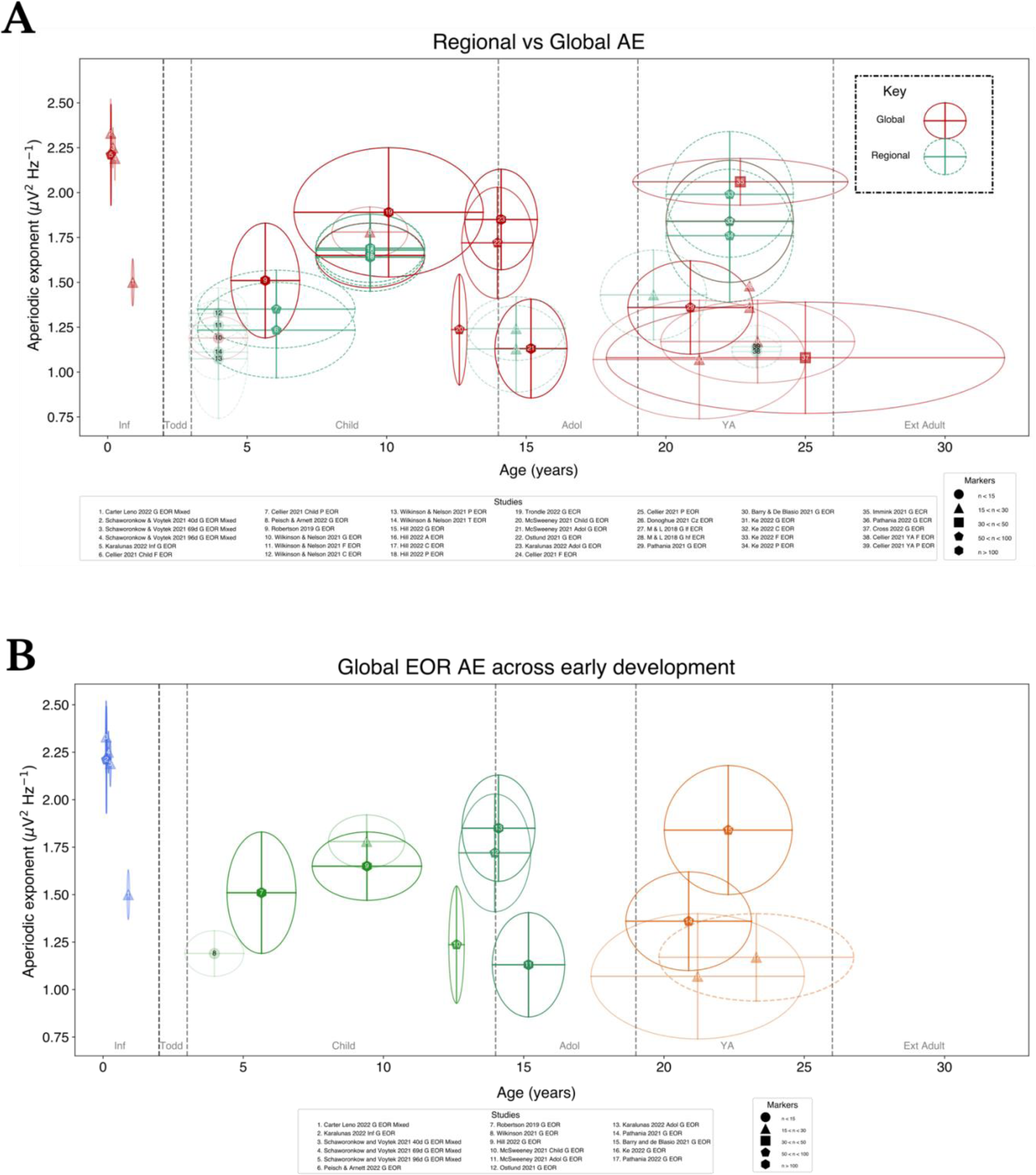
Consistency of the (A) global aperiodic exponent (μV^2^ Hz^−1^) across global and regional scales and (B) focusing explicitly on the global trend of AEs across lifespan stages. For studies where both EOR and ECR were available, only EOR was plotted, as to avoid excess overlap. (A) dotted lines indicate regional AEs, whilst solid lines denote global AEs. Alpha encoding as in Figure 2. Study numbers (white, black) only differ to enhance readability. (B) Colourisation by lifespan stage from infancy-young adulthood. ‘lf’ and ‘hf’ denote low and high frequency slope estimation ranges.

In EOR, we observe age-related AE stabilisation following infancy, with the centre of this trajectory in line with the infant AE estimates of Carter Leno *et al*., (2022)(**Figure 3B**). In summary, there is insufficient evidence in infancy-toddlerhood to validate exponential AE decay, and from childhood onwards AEs seem to vary (partly due to the broad spread of ages in individual studies, as reflected in the age SDs). Whether consistent AE decreases occur from infancy to toddlerhood is likely to be better revealed by studying data at the individual level, dissecting both within- and between-study variability with greater precision. This includes exploring the impact of parameter decisions, such as the number of peaks fit and peak height which affect slope estimation and therefore AE estimates. Notably, the topography of the AE changes with age (**Figure 4**) with AE maxima shifting from posterior foci during infancy (and early toddlerhood) to the midline with continued development. Comparatively, changes in the HE across early development are equally subtle, with evidence from the majority of included studies illustrating that HEs vary by < 0.10 for any given band across the early life span.

**Figure 4.**
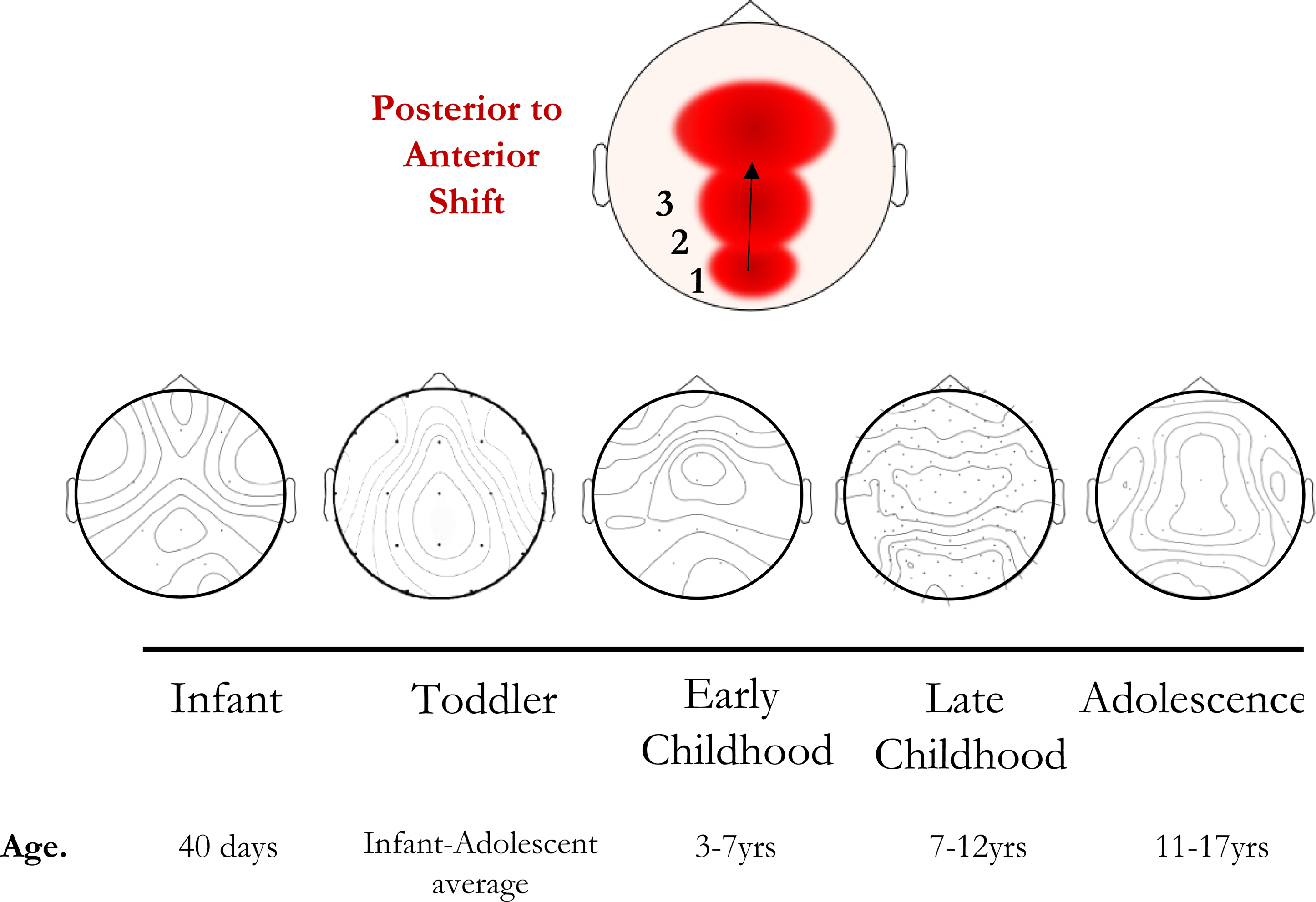
Illustrative regional maturation of the aperiodic exponent (μV^2^ Hz^−1^) with age. Due to limited access to study data for studies in each lifespan stage, topoplots have been generated from available eyes-open rest (EOR) data (references: #1, #9, #11, #15, #2 respectively). For the toddler topoplot, transparency edits to the corresponding published topoplot were made as the data were not publicly available or supplied on request.

### Hurst Exponent: Infancy-Young adulthood

For the HE synthesis, twenty-four studies were included: seven containing infants, one containing toddlers, four containing children, three containing adolescents and nine containing young adults. Infants displayed persistent neural activity patterns (HE>0.50) from delta-beta bands (1-30Hz), during wake (Smith *et al*., 2017) and sleep (Smith *et al*., 2021). Whilst no studies explicitly examined HE maturation in infancy or toddlerhood, one study used a sample containing toddlers. Houtman *et al*., (2021) showed that in an infant-toddler sample, HEs vary from ∼0.65-0.68 across the scalp (with no statistical difference observed between toddlers and children). In childhood, studies collectively showed similar HE in older children, with HE also >0.50: Bruining *et al*., (2020) showed global HE of 0.66±0.04 during ECR in older children (the same sample used in Houtman’s work). Moreover, in a study from ages 5-71yrs (5-7, 16-18yrs longitudinally), Smit *et al*., (2011) identified age-related changes in alpha (5-18yrs occipital maxima, 25yrs: parietal maxima) and beta band HEs (5, 7yrs frontal maxima, 16-50yrs: occipital maxima). By contrast, the theta band HE of Smit *et al*., (2011) are stable and parietal dominant from childhood (5yrs) until YA (25yrs) before switching to frontal dominant in adolescence. Conversely, Kwok *et al*., (2019) observed anti-persistent trends (HE<0.50) for global alpha band HE (EOR: ∼0.09, ECR, ∼0.06) and identified ECR-EOR differences unrelated to age. HEs remain consistent throughout adolescence, with ECR alpha and beta band HE (Linkenkaer-Hansen *et al*., 2007) similar across studies (Gao *et al*., 2017). In YA, Smit, Linkenkaer-Hansen and de Geus (2013) identify EOR alpha band HE maxima in the central midline consistent with other ECR and EOR studies (Linkenkaer-Hansen *et al*., 2001). Moreover, other EOR studies illustrate increasing HE with age; both Nakao *et al*., (2019) and Liu *et al*., (2022) reported consistent ECR alpha band HE across the scalp. However, Natarajan *et al*., (2004) report considerably lower global HE (0.29 vs ≥0.70-0.80 in other studies), and Sleimen-Malkoun *et al*., (2015) identify a broadband global HE of 1.69, suggesting non-stationary (variation unrestricted to a singular mean/setpoint) with occipital maxima and frontal minima. Overall, studies continue to demonstrate posterior (occipital) HE maxima for the alpha band (Irrmischer *et al*., 2018; Bornas *et al*., 2013) consistent with prior power-based studies. In addition, Irrmischer *et al*., (2018) showed that for theta and beta bands, global EOR HE exceed ECR HE. Overall, the HE lifespan trend entails mostly subtle increases in HE with age, differing depending on the band examined and falling within a range of 0.60-0.80 (non-stationary and persistent).

### Aperiodic/Power Law Exponents: Infancy-Young adulthood

For the AE/PLE synthesis, thirty-six studies were included: five containing infants, two containing toddlers, ten containing children, five containing adolescents and fourteen containing young adults. Infant AE (typically >2.00) was higher than in any other lifespan stage for the included studies and appeared to decrease throughout infancy. For instance, Schaworonkow and Voytek (2021) described global AE decreases from 40-134 postnatal days with posterior maxima (40-70days: 3.21, 70-96days: 2.95, 96-134 days: 2.75). In younger infants (0.12 vs 0.81yrs), Karalunas *et al*., (2022) observed EOR-ECR AE averages that were maximal in the midline (2.48, Cz). Conversely, Carter Leno *et al*., (2022) studied 10-month-old infants and identified global AE of 1.50, with no significant regional AE differences nor age effects. By contrast, the evidence from PLE studies shows much lower 1/*f* estimates; during movie-watching, infant global PLE was ∼0.58 for Roche *et al*., (2019), considerably below global PLE observed by Fransson *et al*., (2013) (2.07, occipital cortex).

The largest gap in the developing AE literature sits in toddlerhood; toddler AE are the least characterised of the studied lifespan stages, with only one study evident (Houtman *et al*., 2021), which described AE that were maximal in the midline (∼1.50-1.60). Additional insights were gained from the toddler sub-cohort (N=5) of Cellier *et al*., (2021) wherein steeper posterior (1.27-1.83) than frontal (0.47-1.81) AE were observed. Moreover, Cellier *et al.,* found that AEs significantly decreased with age across their full cohort (3-24yrs) from toddlerhood through young adulthood (*r* = −0.36).

In comparison with other lifespan stages, AE and HE have been best characterised in childhood. Childhood studies recruited TD (McSweeney *et al*., 2023; Tröndle *et al*., 2022; Cellier *et al*., 2021) and case-control child cohorts for comparison with neurodevelopmental conditions including ADHD (Peisch and Arnett, 2022; Arnett, *et al.,* 2022a, b) and Fragile X syndrome (Wilkinson and Nelson, 2021). Childhood AE studies show higher AE than in infancy-toddlerhood, shifting from negative linear AE decay to a positive trend from early to late childhood. Studies in overlapping ages for early (Wilkinson and Nelson, 2021; Robertson *et al*., 2019; Peisch and Arnett, 2022; Hill *et al*., 2022) and late childhood (Tröndle *et al*., 2022; Ostlund *et al*., 2021; McSweeney *et al*., 2021) are generally in agreement in terms of both the direction and range of AE, for both ECR and EOR (**Figure 2B**) and regional versus global (**Figure 3A**) respectively. This was also consistent with figure estimates for studies where data could not be directly obtained (Houtman *et al*., 2021). Two studies provided statistical evidence of a negative age-related AE trend; firstly by Peisch and Arnett (2022) in younger children (*r* =-.30, consistent with previous overlapping work: Arnett *et al.,* 2022a, b), and secondly by McSweeney *et al*., (2023) in older children where a quadratic age-related AE decrease was observed, and ECR AE (1.77) significantly exceeded EOR (1.53) (a trend shown in other studies across the early lifespan, see **Figure 2B**).

Two studies in childhood which have both AE data for ECR and EOR demonstrated these conditions to be comparable (Hill *et al*., 2022; Ostlund *et al*., 2021), alongside single-condition data (typically EOR) from other studies in this stage (**Figure 2B**). Equally, two studies with AE measures for both global and regional scales highlighted comparability across scales (Wilkinson and Nelson, 2021; Hill *et al*., 2022), with similar trajectories evident based on combined data from other studies as in **Figure 3A** (Robertson *et al*., 2019; Ostlund *et al*., 2021; Peisch and Arnett, 2022; Tröndle *et al*., 2022; McSweeney *et al*., 2021). Significant relationships between AE for both scales have also been reported for EOR but not ECR (Hill *et al.,* 2022: [global] *r* = -.24, [regional] anterior: *r* = −0.28, central: −0.24, posterior: −0.35). Topographically, AE maxima in late childhood seem to be parietal dominant (Peisch and Arnett, 2022; Tröndle *et al*., 2022).

In adolescence, three studies provided quantitative evidence for age-related AE decreases (Ostlund *et al*., 2021; McSweeney *et al*., 2021; Karalunas *et al*., 2022), with additional support for this decreasing trajectory in sub-cohort data from Cellier *et al*., (2021). Age-related decreases are observed for both ECR and EOR (Ostlund *et al*., 2021; McSweeney *et al*., 2021), with lower AE observed in females, and faster age-related flattening observed in males (McSweeney *et al*., 2021). Given the collinearity between EOR and ECR, some authors opted to average across conditions (Ostlund *et al*., 2021). Topographic data was only available from one study (Karalunas *et al*., 2022), highlighting AE maxima in the central midline and lateral electrodes (extending more frontally and laterally in higher density caps), with lower adolescent AE than in the study’s infant sample.

A more complex trend is observed during YA, with divergent lines of evidence suggesting an increased versus decreased age effect when taken as a whole. Early YA resting-state PLE studies report considerably lower estimates than studies leveraging methods accommodating for oscillatory peaks to derive AE. For example, Smit, Linkenkaer-Hansen and de Geus (2013) identify EOR PLE maxima in the central midline (0.20-0.40) whilst Muthukumaraswamy and Liley (2018) use IRASA to account for knees in the spectra by modelling multiple slopes, identifying global AE of 1.36 (β_1_:0.1-2.5Hz, frontal maxima) and 1.48 (β_2_:20-100Hz, central maxima) respectively. AE studies cluster between 1.30-1.60, similar to the range described by Merkin *et al*., (2023), irrespective of whether FOOOF (Donoghue *et al*., 2020; Pathania *et al*., 2021, 2022; Zsido *et al*., 2022; Cross *et al*., 2022), IRASA (Muthukumaraswamy and Liley, 2018) or other methods (Barry and De Blasio, 2021) are utilised, and with similar patterns for ECR and EOR, though ECR AE remains higher. Two exceptions to this are noted (Ke *et al*., 2022; Immink *et al*., 2021), with one of these (Immink *et al*., 2021) identifying ECR AE estimates falling within the tentative infant AE range (>2.00). Merkin *et al.,* also noted that regional (but not global) age-related AE changes were significant when accounting for peak parameters and goodness of fit and did not differ by region.

In YA, the magnitude of ECR AE exceed that of EOR AE (Pathania *et al*., 2022; Cross *et al*., 2022; Barry and De Blasio, 2021), and topographical maxima centre around the central and frontal regions (Pathania *et al*., 2022; Barry and De Blasio, 2021), with an indication that this is more commonly frontal dominant (Ke *et al*., 2022). The differences in regional AE are smaller than in other lifespan stages, thus differences between these maxima (*e.g.* parietal - Pathania *et al*., 2021 vs occipital - Pathania *et al*., 2022) are unlikely to reflect biological differences.

## Discussion

In this systematic review, we aimed to explore how and when EEG derived 1/*f* measures change in early human development, and where variability within early lifespan stages exists. We found that AE and HE age-related changes have complex developmental patterns; (1) HE consistently exceeded 0.50 across development, suggesting persistent and non-stationary signals throughout the early lifespan (2) provisional evidence suggests AEs decrease throughout infancy (*i.e.* an increased excitation:inhibition ratio) prior to the AE varying within confined ranges across subsequent development, (3) this pattern is generally consistent across AE methods, (4) the magnitude of ECR AEs exceed that of EOR AEs throughout early development (with overlapping trends observed), (5) heterogenous post-infancy AE changes do not differ between global or regional scales and (6) a posterior-anterior shift in maximal AE occurs from infancy through young adulthood.

### Further evidence is required to determine age-related AE trends

Despite the influence of narrowband oscillations on slope fitting and exponent estimation, PLEs show age-related decreases (Waschke, Wöstmann and Obleser, 2017). We find that AE changes non-linearly from childhood onwards, a finding in line with large child AE datasets in both the EEG (Cellier *et al*., 2021; McSweeney *et al*., 2023) and MEG (Thuwal, Banerjee and Roy, 2021) literature. However, several other EEG studies fail to identify global (Merkin *et al*., 2023) or regional age-effects (Hill *et al*., 2022). As recent evidence suggests that the balance of E:I in early infancy may have key implications for brain development and function across the lifespan, infant AEs can provide an important early non-invasive marker of the integrity of functional brain activity. Provisional evidence shows decreases in global AEs from the first several weeks after birth in term-born infants, however, there are significant gaps in the literature, particularly in mid and late infancy. Recently, Rico-Picó *et al*., (2023) identified early decreases in global AE (6-9 months) and flattening thereafter (9-18 months). In toddlerhood, gaps in characterising AE are more significant, which impedes the interpretation of a qualitative ‘trajectory’ of AE development thereafter (particularly given the complex patterns of AE variability observed in childhood). A preprint by Wilkinson *et al*., (2023) partially addresses this toddler AE gap, charting resting AE from 2-44 months, highlighting considerably flatter spectra than we observe here, with AE rising from 1.00 to 1.20 (0-1200 days) and age-sex interactions being evident. Evidence from this work suggests that AE increases persist through infancy and toddlerhood. Physiologically, increased postnatal AEs (higher inhibition/lower excitation) are in keeping with the axiom that postnatal GABA-related activity shifts from excitatory (depolarising) to inhibitory (hyperpolarising) postnatally due to changes in intracellular chloride concentrations (Ben-Ari *et al*., 2007; Kirmse *et al*., 2015; Ben-Ari and Cherubini, 2022). However, as this GABA shift occurs in immature neurons, E:I balance later tilts towards excitation (during or following late infancy) as circuits and networks mature and glutamate signalling predominates. Whether net excitation or inhibition initially dominates activity in the developing brain is frequently disputed: evidence from rodent (postnatal days 2-12) and newborn infant frontal EEG (35-46 postmenstrual weeks) 1/*f* data show higher AEs are observed with increasing age (Chini, Pfeffer and Hanganu-Opatz, 2022), possibly due to a more protracted integration of interneurons (relative to pyramidal neurons) into emerging circuits.

Furthermore, there is currently no consensus on whether at the earliest point in infant development E:I balance tilts more towards excitation or inhibition, as AE measure coverage in this window is limited. Wilkinson *et al* (2023)’s data suggests that all regional AEs but temporal AEs increase during infancy, whilst temporal AE decreases prior to a nadir∼400 days, before increasing. Overall, these data and our findings agree that infant AE maxima are in the posterior channels overlaying occipital regions. However, our findings differ as to the direction of expected regional/global AE change. Whilst higher frequency peaks in Wilkinson’s data may have affected AE estimation, the authors performed comprehensive model fit screening. They modified slope fitting functions in order to accommodate poor estimations at 10-20Hz, whereas modelling of knees or multiple slopes (as in Shuffrey *et al*., 2022) such as from 0.5-10Hz may have produced different findings. It is unclear whether this point could also extend to Chini *et al’s* data from early infancy. Overall, future AE research should seek to provide robust estimates of early neonatal AEs, identifying whether age-related decreases occur from the beginning of the neonatal period and continue through to late infancy. One approach to address this involves visualising pooled individual-level data across studies to get an accurate consensus of how AE varies within given stages (*e.g.* infancy) and, consequently, how variable it becomes across lifespan stages into adulthood. Importantly, the results of the group-level analysis reported here suggest that such an analysis should consider differing methodologies and model-fitting parameters to make robust comparisons.

### Comparable AE results across methods

In contrast to early infancy, synthesising AE patterns across methods in subsequent childhood suggest that AE estimation methods are broadly comparable (except for converted HE): in YA in particular, FOOOF, IRASA, PawWNextra and PLE estimates overlap from 0.77-1.93. HEs differ in that whilst they inform us about the temporal persistence of EEG activity patterns (revealing when patterns become conserved across time), HEs are agnostic to the direction of change and tilting of the E:I balance spectra and thus only partially explain how E:I shapes evolving functional circuits/networks. Moreover, converted HE show substantial differences vs AE measured directly, likely as a result of: (1) many HE being characterised via DFA based on amplitude envelopes of specific bands (particularly papers from >2012), (2) time domain effects occurring due to reduced sampling windows and recording lengths, (3) data self-similarity which must be verified in source data for DFA and (4) conversion assuming DFA scaling exponents (⍺) are Gaussian (*i.e.* >0.5 or < 0.5) and not Brownian (∼0.5)(Eke *et al*., 2002). Finally, (5) the HE does not accommodate for oscillatory influence, and thus similar to PLEs, converted AE will provide potential over- or underestimates of the true AE, potentially providing physiologically implausible estimates akin to what has been described in the frequency spectra literature (Barry and De Blasio, 2021). Across the lifespan, HE studies consistently show persistent (0.50<HE<1.00), non-stationary (HE>0.50) patterns in each lifespan stage, reminiscent of sustained processing during measurement, and notably, of properties of a system with memory whose signal is exhibiting positive correlations over time (Hardstone *et al*., 2012). Only Sleimen-Malkoun *et al*., (2015)’s study suggested non-stationarity (1>⍺>2, therefore HE=⍺-1) in the EEG during rest. It is, however, worth noting that detecting developmental changes in HEs requires both large subject samples and (noise-free) long epochs (Berthouze, James and Farmer, 2010) in order to characterise temporal correlations at multiple scales. This is all the more poignant given the individual variation in long-range temporal correlations within and across subjects (Linkenkaer-Hansen *et al*., 2007). Whilst the HE has been studied more deeply in adults, larger AE studies tend to focus on childhood. The HE studies captured by this systematic review were generally monofractal or equivalent multifractal measures (H(2)), though recent studies have tended to focus on multifractal EEG dynamics, potentially offering insights into more complex non-stationary scaling behaviours in EEG data (Zorick and Mandelkern, 2013); these methods thereby index E:I proxies at multiple temporal scales, akin to estimating 1/*f* slopes across multiple frequency ranges.

### ECR consistently exceeds EOR AE

When focusing on AEs specifically, there was consistent evidence that the magnitude of ECR AEs exceeded that of EOR AEs across the lifespan. Childhood studies showed marginal AE increases in later (Hill *et al*., 2022; Tröndle *et al*., 2022; McSweeney *et al*., 2021) versus earlier (Robertson *et al*., 2019; Wilkinson and Nelson, 2021) childhood (for both EOR/ECR), although this may reflect greater between-dataset variability rather than genuine AE increases preceding a prolonged age-related decline. Across early development, the greater magnitude of ECR vs EOR AE is driven not only by posterior dominant alpha band activity (Wilson, Castanheira and Baillet, 2022) but also the activity in other frequency bands (Barry *et al*., 2007). Whilst some authors using other methods report FOOOF AEs are greater in EOR than in ECR, such as SPRiNT (Wilson, Castanheira and Baillet, 2022), our findings consistently show that studies using FOOOF, IRASA and PaWNextra find ECR AE to exceed that of EOR. Moreover, both EOR and ECR AEs follow similar trajectories suggesting these ‘resting’ E:I processes mature in similar ways, consistent across both regional and global scales.

### Regional vs Global AE

For the most part, global AE magnitude exceeds that of regional AEs (**Figure 3A**), likely owing to the average rate of AE decrease across the scalp remaining constant across development whilst regional AE differs (as maturing regions shift developmentally). For example, regional changes in AE are apparent in early childhood, but equilibrate before YA and thus the neurobiological changes underlying AE changes during mature ageing are likely physiologically distinct from those in the earlier lifespan (Merkin *et al*., 2023). For example, using simultaneous EEG-fMRI during EOR, Jacob *et al*., (2021) identified posterior parietal AE maxima (∼1.60) in adults, with global average AE (1.49) being associated with decreases in frontal and increases in cerebellar, insular and cingulate blood-oxygen-level-dependent fMRI activity. In later life, Aggarwal and Ray (2023) identify there are no age-related MEG AE differences in younger versus older adults up to 50Hz, but from 64-140Hz AE decreases and from 230-430Hz increases (higher inhibition), collectively suggesting that subtle GABAergic changes may occur in later life outside of spectral ranges accessible to EEG. Moreover, whilst AE maturation may taper in earlier development, AE development is not static thereafter. Differences between mid and older adults were evident in posterior channels, similar to what is observed in early development, suggesting that network hubs established in infancy are also the last to change during later ageing. However, it is worth considering that when producing regional estimates to inform network maturation, selecting spatially neighbouring high SNR channels is vital (Linkenkaer-Hansen *et al.,* 2001). Therefore, whether estimating regionally or globally, researchers should utilise model fit statistics to ensure adequate representation of underlying neural data. Currently, only a minority of studies report model fits and fewer still include fits as covariates. Given the need to systematically validate lifespan AE, we consider model reporting to be vital to ensuring accurate characterisation of developmental trajectories.

### Regional AE maxima shift across typical development

Topographical E:I maxima by definition relate to spectra with lower E:I balance (steeper AE spectra) relative to the rest of the brain, which in the absence of a stimulus (endogenous or exogenous) may suggest ongoing regional maturation (as opposed to flatter spectra and greater neural “noise” in ageing and pathology (Dave, Brothers and Swaab, 2018; Pertermann *et al*., 2019), or regions involved in networks which are more selectively held at baseline during conditions of rest. Understanding where AE are maximal (and neural noise minimal) provides insights into which regions are potentially undergoing maturational changes, which must first be characterised in TD to provide a referential maturational trajectory. We find that typical AE maturation displays a posterior-to-anterior shift in ageing (PASA), similar to the fMRI literature, with age-related reductions in occipital activity concomitant with increasing frontal activity (Davis *et al*., 2008; McCarthy, Benuskova and Franz, 2014). In longitudinal infant data at 6, 9 and 16 months, Rico-Picó *et al*., (2023) show AE decreases more slowly in occipital (maximal) and frontal versus parietal and central areas. These AE changes temporally coincide with white matter maturation and ongoing activity integration in the toddler as sensorimotor skills emerge (Hagmann *et al*., 2010). From childhood to late adolescence, the difference between posterior and anterior AE seems to grow with age across both sleep and wake, being the strongest in the second stage of sleep (Favaro *et al*., 2023). fMRI FC maturation at this point in development follows a sensorimotor-association gradient where primary sensory maturation precedes that of frontal executive and association areas (Sydnor *et al*., 2023). Overall, AEs demonstrate PASA in line with prior neuroimaging evidence, and accordingly, centro-frontal regions seem to be the maturational ‘endpoint’ for early AE development (with the lowest neural noise), perhaps supporting an increasing processing requirement for cognitive function.

### Limitations and future work

This review highlights the complexity of characterising group-level age-related AE changes across studies, methods, and spatial scales. Given the heterogeneity in AE estimates, AE must be estimated on relatively noise-free data (minimal evidence of physiological/non-physiological artefacts including eye movements, electrode bridging, line noise, cardiac and respiratory signals or sweat-induced artefacts). Simulation work suggests SNR>2 are appropriate for determining HEs (Linkenkaer-Hansen *et al*., 2007). SNR may also be influenced by equipment selection, particularly for acquisitions with reduced channels, poorer contact quality and/or more flexible sensors, designed for ‘active’ paradigms (see Grummett *et al*., 2015). There are further influences due to data constraints and processing decisions, including the effect of window length on smoothing, affecting peak estimates. Data reference schemes will also affect PSD and AE estimates (Gao, Peterson and Voytek, 2017), for which most included studies used average referencing (see **Table 1** and **Supplementary Material IV**). Equally, filtering decisions affect the frequency range available for exponent estimation; as others have shown, estimations on lower versus higher frequency slopes differ (Shuffrey *et al*., 2022; Muthukumaraswamy and Liley, 2018) and may have different physiological interpretations in the contexts of neurodevelopment and pathophysiology. In addition, motion is generally unavoidable in infants and children, who may not tolerate prolonged recording periods, hindering attempts to use epoch averaging to increase SNR.

AE estimation methods must, therefore, be valid for the applied dataset(s) and comparable with prior studies. For instance, when comparing FOOOF and IRASA results, a consideration is that IRASA evaluates spectral ranges beyond the fitted range in order to compute median AE using resampling factors (Gerster *et al*., 2022). Therefore, comparing results by exact frequency mapping results in evaluating upper or lower limit ranges which may be affected by filtering or noise, thus biasing AE estimation. Fortunately, IRASA and FOOOF AE estimates in this review overlap heavily, but this is nonetheless a consideration. Moreover, Gyurkovics *et al*., (2021) highlight that neural variability as captured by the 1/*f*^β^ may differ between age groups, and spectra calculated from longer epochs (or averages) may be optimal for FOOOF as shorter single-trial spectrum models can overfit noise (due to the number of free parameters). For specific limitations and strengths of either method, see Gerster *et al*., (2022). Other tools for parameterising the AE have been introduced recently, but these have yet to be applied to resting scalp EEG (Seymour *et al*., 2022). Beyond methodological choices, AEs may also change based on genotype, task paradigm, and cognitive state (He, 2014; Voytek *et al*., 2015; Donoghue *et al*., 2020), hence the focus on quasi-resting states and typical development in this review. The reviewed literature discussed provided (predominantly) cross-sectional measures of AE across development. However, the AE varies both statically across (*e.g.* 0.68-2.77 in Tröndle *et al*., 2022) and dynamically within individuals (during recordings), especially in task-specific contexts. For example, Wilson, Castanheira and Baillet (2022) show that the AE varies over time in the resting-state, with YA AEs predictive of subject state (EOR vs ECR). Ultimately, more complex modelling may be required to evaluate how AE variation within-individual differs from results observed across individuals. Moreover, we suggest that the pooling of individual datapoints from constituent studies and a quantitative analysis thereof may better distinguish how age (and sex) influence biological AE changes across the lifespan.

In addition to limitations inherent to the methods of studies included in the review, there are limitations to the review itself. Given the sparsity of effect size measures for AE and age (age*AE correlations or age-related mean group differences between EOR and ECR AE) it was not possible to produce a meaningful meta-analytic measure of age-related AE change. In addition, our inclusive search approach (including searches for terms relating to fractal measures to ensure sufficient HE data was sourced to provide adequate converted comparisons) resulted in significant heterogeneity. The suitability of including infant AEs where participant attention was captured using “toys” (Carter Leno *et al*., 2022) as a “resting” AE measure could be disputed. However, we perceive this to be a necessary means to engage young infant participants and minimise motion, and in the trajectories we have qualitatively illustrated, these AE estimates are consistent with a trend of decreasing AEs from infancy towards childhood.

### Summary

In summary, this review demonstrates that age-related AE changes in early development are complex. However, there are significant gaps in the data which currently prevent the robust establishment of age-related directions of change and reliable AE ranges, particularly in infancy and toddlerhood. We identify consistent AEs across methods and scales and confirm higher values of ECR than EOR, as well as developmental changes in AE maxima. However, our review exclusively characterises the maturation of AEs in the resting state. Thus, specific task-related AE changes across the lifespan remain to be explored. As AE data is made available to the community, we can collectively extend the findings of this review to advance knowledge of how E:I shapes FC in early development. Moreover, characterising typical AE development provides a point of reference for exploring atypical development in which early life E:I balance is perturbed, where AEs could act as a potential non-invasive biomarker.

## Acknowledgements

The authors would like to thank those contributing additional data and/or clarifications through personal correspondence, without which our study would not be possible: Anne Arnett, Robert Barry, Virginia Carter-Leno, Dillan Cellier, Zachariah Cross, Thomas Donoghue, Peter Fransson, Maarten Immink, Sarah Karalunas, Keith Lohse, Marco McSweeney, Brendan Ostlund, Natalie Schaworonkow, Dirk Smit, Rachel Smith, Marius Tröndle, Sampsa Vanhatalo, Bradley Voytek and Carol Wilkinson.

## Availability of data, code, and other materials

Data for this review were obtained on reasonable request from corresponding authors and from online repositories, and thus are not within our right to share. For datasets used, see **Table 1**. Visualisation functions scripted to produce **Figures 2 and 3** have been made <u>available</u>.

---

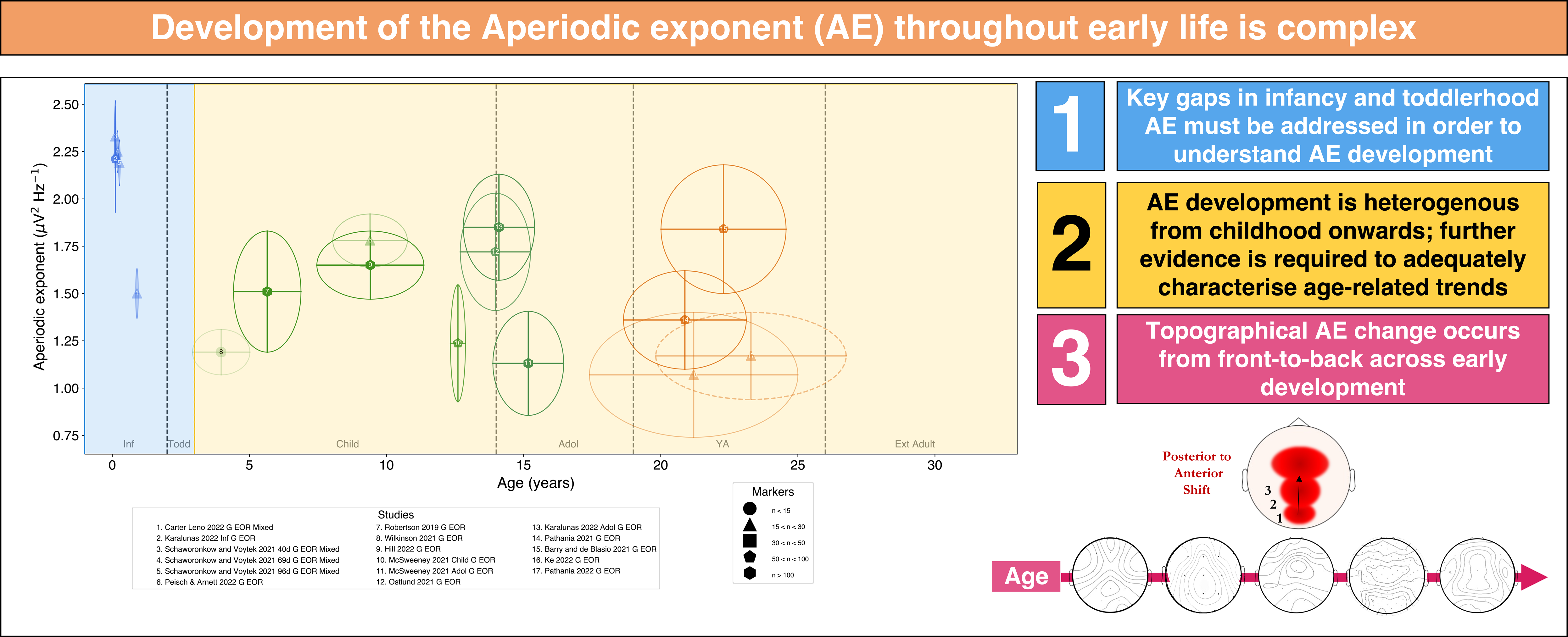

