## Supplementary_Materials for "Aperiodic and Hurst EEG exponents across early human brain development: a systematic review"

***Supplement I. Systematic Review Example Search Strategy***

**Scopus Search Strategy, March 2023:**

TITLE-ABS-KEY ( aperiodic AND ( exponent* OR slope ) ) OR TITLE-ABS-KEY ( aperiodic W/2 ( exponent* OR slope ) ) OR TITLE-ABS-KEY ( hurst AND exponent ) OR TITLE-ABS-KEY ( hurst W/2 exponent ) OR TITLE-ABS-KEY ( ( detrend* AND fluctuation AND analysis ) OR fractal ) AND TITLE-ABS-KEY ( eeg OR electroencephal* ) AND TITLE-ABS-KEY ( birth OR newborn OR neonat* OR infan* OR toddler* OR child* OR teenage* OR adoles cent* OR ( young AND adult ) OR development* OR ( early AND life ) OR maturation* )

***Supplement II. QuADS Risk of Bias Criteria***

| QuADS Criteria | 0 | 1 | 2 | 3 |
|---|---|---|---|---|
| 1. **Theoretical or conceptual underpinning to the research** | No mention at all. | General reference to broad theories or concepts that frame the study. e.g. key concepts were identified in the introduction section. | Identification of specific theories or concepts that frame the study and how these informed the work undertaken. e.g. key concepts were identified in the introduction section and applied to the study. | Explicit discussion of the theories or concepts that inform the study, with application of the theory or concept evident through the design, materials and outcomes explored. e.g. key concepts were identified in the introduction section and the application apparent in each element of the study design. |
| **2. Statement of research aim/s** | No mention at all. | Reference to what the sought to achieve embedded within the report but no explicit aims statement. | Aims statement made but may only appear in the abstract or be lacking detail. | Explicit and detailed statement of aim/s in the main body of report. |
| **3. Clear description of research setting and target population** | No mention at all. | General description of research area but not of the specific research environment e.g. 'in primary care.' | Description of research setting is made but is lacking detail e.g. 'in primary care practices in region [x]'. | Specific description of the research setting and target population of study e.g. 'nurses and doctors from GP practices in [x] part of [x] city in [x] country.' |
| **4. The study design is appropriate to address the stated research aim/s** | No research aim/s stated or the design is entirely unsuitable e.g. a Y/N item survey for a study seeking to undertake exploratory work of lived experiences. . | The study design can only address some aspects of the stated research aim/s e.g. use of focus groups to capture data regarding the frequency and experience of a disease. | The study design can address the stated research aim/s but there is a more suitable alternative that could have been used or used in addition e.g. addition of a qualitative or quantitative component could strengthen the design. | The study design selected appears to be the most suitable approach to attempt to answer the stated research aim/s. |
| **5. Appropriate sampling to address the research aim/s** | No mention of the sampling approach. | Evidence of consideration of the sample required e.g. the sample characteristics are described and appear appropriate to address the research aim/s. | Evidence of consideration of sample required to address the aim. e.g. the sample characteristics are described with reference to the aim/s. | Detailed evidence of consideration of the sample required to address the research aim/s. e.g. sample size calculation or discussion of an iterative sampling process with reference to the research aims or the case selected for study. |
| **6. Rationale for choice of data collection tool/s** | No mention of rationale for data collection tool used. | Very limited explanation for choice of data collection tool/s. e.g. based on availability of tool. | Basic explanation of rationale for choice of data collection tool/s. e.g. based on use in a prior similar study. | Detailed explanation of rationale for choice of data collection tool/s. e.g. relevance to the study aim/s, codesigned with the target population or assessments of tool quality. |
| **7. The format and content of data collection tool is appropriate to address the stated research aim/s** | No research aim/s stated and/or data collection tool not detailed. | Structure and/or content of tool/s suitable to address some aspects of the research aim/s or to address the aim/s superficially e.g. single item response that is very general or an open-response item to capture content which requires probing. | Structure and/or content of tool/s allow for data to be gathered broadly addressing the stated aim/s but could benefit from refinement. e.g. the framing of survey or interview questions are too broad or focused to one element of the research aim/s. | Structure and content of tool/s allow for detailed data to be gathered around all relevant issues required to address the stated research aim/s. |
| **8. Description of data collection procedure** | No mention of the data collection procedure. | Basic and brief outline of data collection procedure e.g. 'using a questionnaire distributed to staff'. | States each stage of data collection procedure but with limited detail or states some stages in detail but omits others e.g. the recruitment process is mentioned but lacks important details. | Detailed description of each stage of the data collection procedure, including when, where and how data was gathered such that the procedure could be replicated. |
| **9. Recruitment data provided** | No mention of recruitment data. | Minimal and basic recruitment data e.g. number of people invited who agreed to take part. | Some recruitment data but not a complete account e.g. number of people who were invited and agreed. | Complete data allowing for full picture of recruitment outcomes e.g. number of people approached, recruited, and who completed with attrition data explained where relevant. |
| **10. Justification for analytic method selected** | No mention of the rationale for the analytic method chosen. | Very limited justification for choice of analytic method selected. e.g. previous use by the research team. | Basic justification for choice of analytic method selected e.g. method used in prior similar research. | Detailed justification for choice of analytic method selected e.g. relevance to the study aim/s or comment around of the strengths of the method selected. |

| | | | | |
|---|---|---|---|---|
| **11. The method of analysis was appropriate to answer the research aim/s** | No mention at all. | Method of analysis can only address the research aim/s basically or broadly. | Method of analysis can address the research aim/s but there is a more suitable alternative that could have been used or used in addition to offer a stronger analysis. | Method of analysis selected is the most suitable approach to attempt answer the research aim/s in detail e.g. for qualitative interpretative phenomenological analysis might be considered preferable for experiences vs. content analysis to elicit frequency of occurrence of events. |
| **12. Evidence that the research stakeholders have been considered in research design or conduct.** | No mention at all. | Consideration of some the research stakeholders e.g. use of pilot study with target sample but no stakeholder involvement in planning stages of study design. | Evidence of stakeholder input informing the research. e.g. use of pilot study with feedback influencing the study design/conduct or reference to a project reference group established to guide the research. | Substantial consultation with stakeholders identifiable in planning of study design and in preliminary work e.g. consultation in the conceptualisation of the research, a project advisory group or evidence of stakeholder input informing the work. |
| **13. Strengths and limitations critically discussed** | No mention at all. | Very limited mention of strengths and limitations with omissions of many key issues. e.g. one or two strengths/limitations mentioned with limited detail. | Discussion of some of the key strengths and weaknesses of the study but not complete. e.g. several strengths/limitations explored but with notable omissions or lack of depth of explanation. | Thorough discussion of strengths and limitations of all aspects of study including design, methods, data collection tools, sample & analytic approach. |

### *Supplement III. Risk of bias assessment for included studies*

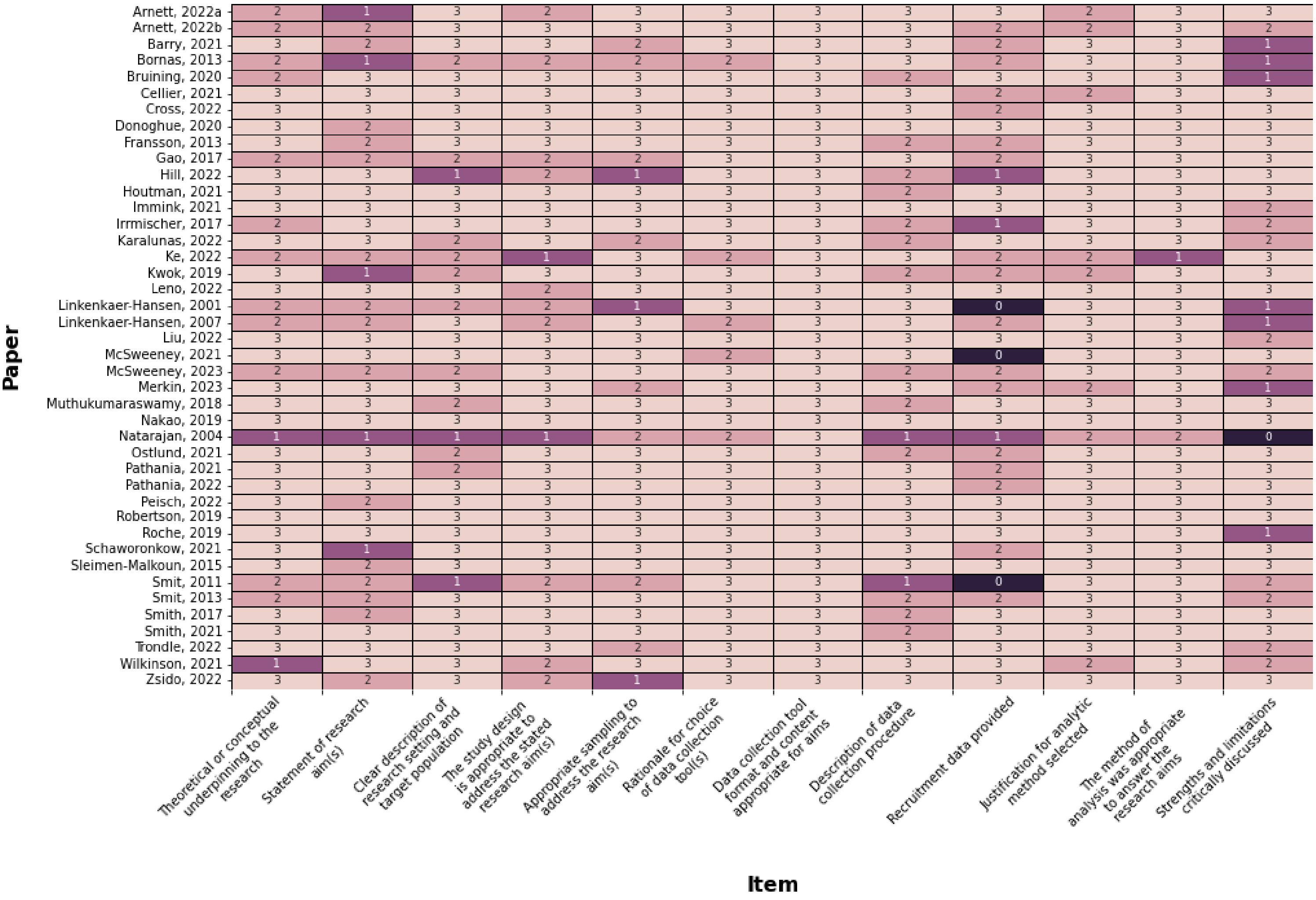

***Supplement IV.*** **Studies included in the review, with technical details**

An extension of **Table 1**. (main text) with additional technical details regarding recording length, referencing, models and pertinent temporal windows for measure calculation (PSD estimation/sliding windows).

| # | Study | Lifespan Stage (age, yrs) | Measure | Technical Specs | Scale(s) | Original Measure | HE to AE | Measure | N (M, F) | Source | F |
|---|---|---|---|---|---|---|---|---|---|---|---|
| 1 | Schaworonkow & Voytek (2021) | Infancy<br>(0.10-0.56) | 1/f<br>(FOOOF) | RECORDING, EPOCH<br>5 mins; 10s epoch length;<br>PSD ESTIMATION<br>Multitaper<br>REFERENCING<br>Offline: Common average<br>MODEL<br>1-10Hz; peak_width_limits=[0.5, 12.0], maximum_n_peaks=5, min_peak_height=0, peak_threshold=2.0, aperiodic_mode='fixed'. | Channelwise | S1: 1.74-3.22 (N = 20)<br>S2: 1.74-2.95 (N = 20)<br>S3: 1.79-2.25 (N = 20)<br>S4: 1.94-2.98 (N = 5)<br>S5: 1.46-2.76 (N = 3)<br>S6: 1.88-2.63 (N = 2) | | Baseline-wakeful reaching | 22(10,12) | Methods,<br>Auth Corr.,<br>GitHub | |
| 2 | Karalunas (2022) | Infancy<br>(0.12±0.01)<br><br>Adolescent<br>(14.10±1.30) | 1/f<br>(FOOOF) | RECORDING, EPOCH<br>8 mins; 2s epoch length<br>PSD ESTIMATION<br>Hanning window; no further information given<br>REFERENCING<br>Offline: Common average<br>MODEL<br>Infant 1-30Hz, Adolescent 2-50Hz<br>peak_width_limits=[1,8],<br>maximum_n_peaks=6, min_peak_height=0,<br>peak_threshold=2.0, aperiodic_mode='fixed'. | Global,<br><br><br>Channelwise | Infant EOR (PEACH cohort): 2.21±0.28<br>Adol EOR (1.85±0.28): ECR (1.98±0.26), EOR-ECR avg (1.91±0.28)<br>Infant: 2.48±0.24 (Cz, EOR)<br>Adol: 2.28±0.19 (Cz, EOR), 2.33±0.27 (Pz, ECR), 2.29±0.19(EOR-ECR average) | | EOR, ECR<br>EOR-ECRavg | 69 (36,33)<br>152(85,67) | Auth Corr. | |
| 3 | Fransson (2013) | Infancy<br>(0.81, 0.75-0.85) | 1/f (PLE) | RECORDING, EPOCH<br>Unknown; 5 mins<br>PSD ESTIMATION, REFERENCING<br>Unknown<br>MODEL<br>0.2-30Hz | Global,<br>Regional,<br>Channelwise | 2.07±0.22 | | Natural Active/ Quiet Sleep | 15(9,12) | Fig 4 | Y |
| 4 | Carter-Leno (2022) | Infancy<br>(0.90±0.05) | 1/f<br>(FOOOF) | RECORDING, EPOCH<br>3 mins social, 3 mins non-social; 1s segment<br>PSD ESTIMATION<br>FFT; 1Hz bins<br>REFERENCING<br>Average<br>MODEL<br>1-10Hz. peak_width_limits=[2,8], maximum_n_peaks=4, peak_threshold=0.1, aperiodic_mode='fixed'. | Global,<br>Regional,<br>Channelwise | 1.50±0.13 (non-social), 1.52±0.16<br>Fz: social (1.53±0.16), non-social (1.51±0.13)<br>Cz: social (1.51±0.15), non-social (1.49±0.13)<br>Pz: social (1.51±0.16), non-social (1.49±0.13) | | ~EOR (social and non-social videos) | 24(13,11) | Table 1,<br>Fig 4,<br>Auth Corr. | |
| 5 | Roche (2019) | Infancy<br>(1.92-10.25) | 1/f (PLE) | RECORDING, EPOCH<br>5-10 mins; 1s, non-overlapping<br>PSD ESTIMATION<br>FFT, Hanning window.<br>REFERENCING | Global$^{est}$,<br>Regional | ~0.58 | | ~EOR (movie) | 37(0,37) | Methods,<br>Results | |

| | | | | | | | | | | |
|---|---|---|---|---|---|---|---|---|---|---|
| | | | | Online: vertex (Cz), Offline: Common average<br>MODEL<br>2-24Hz | | | | | | |
| 6 | Smith, R. (2021) | Infancy<br>(med. 0.63, 0.43-0.82) | HE | RECORDING, EPOCH<br>~19h; > 2 wake, 2 sleep epochs;<br>REFERENCING<br>Linked ear<br>HURST SCALES<br>Box sizes 1- 1/10th signal length (max 120s) | Global | Delta[1-3Hz]: ~0.80 (A), 0.68(S)<br>Theta[4-7Hz]: ~0.74(A), 0.68(S)<br>Alpha[8-12Hz]: ~0.69(A), 0.68(S)<br>Beta[13-30Hz]: ~0.88(A), 0.72(S) | | Awake, Sleep | 20(12,8) | Section 3.1,<br>Fig 6,<br>Auth Corr. |
| 7 | Smith, R. (2017) | Infancy<br>(med. 0.58, 0.48-0.94) | HE | RECORDING, EPOCH<br>Unknown<br>PSD ESTIMATION<br>50% window overlap<br>REFERENCING<br>Unknown<br>HURST SCALES<br>Log(3-25s) windows | Global [s] | Delta[1-3Hz]: ~0.78<br>Theta[4-7Hz]: ~0.70<br>Alpha[8-12Hz]: ~0.66<br>Beta[13-30Hz]: ~0.94 | | Awake<br>~(EOR) | 21(?,?) | Fig 5,<br>Auth Corr. |
| 8 | Cellier (2021) | Toddler (N=5),<br>Child (N=81),<br>Adolescent (N=22),<br>Young Adult (N=8)<br>(2.95-24) | 1/f<br>(FOOOF) | RECORDING, EPOCH<br>Unknown length, 512ms<br>PSD ESTIMATION<br>Welch, 45% overlap, 512 and 1024ms sliding windows<br>REFERENCING<br>Average<br>MODEL<br>1-40Hz. peak_width_limits=[1, $2_{MIPDB}$ /$4_{SRS}$], maximum_n_peaks=4, peak_threshold=2.0, aperiodic_mode='fixed'. | Regional<br>(Parietal-midline [P],<br>Frontal-midline [F]) | Toddler: [P] 1.45±0.23, [F] 1.32±0.54<br>Children: [P] 1.23±0.25, [F] 1.34±0.22<br>Adolescents: [P] 1.24±0.18, [F] 1.13±0.24<br>Young Adults: [P] 1.14±0.12, [F] 1.11±0.09 | | EOR | 116<br>(33,24,59 unlabelled) | Fig 2,<br>Sections 2.2, 3.1,<br>Auth Corr.,<br>OSF |
| 9 | Houtman (2021) | Toddler (2.92 [N=8], 3.92 [N=13]),<br>Child<br>(7-16[N=29]) | 1/f<br>(FOOOF)<br>, HE | RECORDING, EPOCH<br>1-19 min length (multi-cohort); 1s epoch length;<br>PSD ESTIMATION<br>Welch, 2s Hamming, 50% overlap<br>REFERENCING<br>mastoid reference (1/2 TD studies)<br>MODEL<br>1-30Hz; peak_width_limits = [1,6],<br>maximum_n_peaks = 6,<br>min_peak_height = 0.05,<br>peak_threshold = 1.5,<br>aperiodic_mode = "fixed"<br>HURST SCALES<br>log(4)-log(20) (< 8Hz), log(2)-log(20) (>8Hz) | Global<br><br>Channelwise | Infant-toddler (I) & child-adol (C): HE, 11-18Hz: I: ~0.655, C: ~0.656<br>Infant-toddler (I) & child-adol (C): AE, ~1.11-1.60<br>(Hurst, 11-18Hz): I: ~0.63-.70), C: ~0.64 -0.74, 0.66±0.02 | | EOR | 50 (28,22):<br>Inf-Todd:<br>21 (14,7)<br>Child-Adol:<br>29 (14,15) [a] | Fig 3, 5<br>Supp. Fig 5 |
| 10 | Wilkinson & Nelson (2021) | Child<br>(3.98±1.09, 2.67-6.67) | 1/f<br>(FOOOF) | RECORDING, EPOCH<br>2-5 mins; 2s segment<br>PSD ESTIMATION | Global<br>Regional | 1.19±0.12<br>Frontal: 1.26±0.13<br>Central: 1.33±0.14 | | EOR | 12(12,0) | Methods,<br>Results,<br>Auth Corr. |

| | | | | | | | | | | |
|---|---|---|---|---|---|---|---|---|---|---|
| | | | | Multi-tapering, 4s window, 1s step size<br>REFERENCING<br>Online: Cz; Offline: average.<br>MODEL<br>2-55Hz. peak_width_limits=[1, 18], maximum_n_peaks=7, peak_threshold=2.0, aperiodic_mode='fixed'. | | Temporal: 1.11±0.15<br>Posterior: 1.07±0.32 | | | | |
| 11 | Robertson (2019) | Child (5.65±1.23) | 1/f (FOOOF) | RECORDING, EPOCH<br>1-2 min length; 1s epoch length;<br>PSD ESTIMATION<br>Welch, 1s Hamming, 50% overlap<br>REFERENCING<br>Common average<br>MODEL<br>4-50Hz. peak_width_limits=[1, 8], maximum_n_peaks=8, peak_threshold=2.0, aperiodic_mode='fixed'. | Global Channelwise | 1.51±0.32 | | EOR | 50(36,14) | Table 1, Fig 2A, B |
| 12 | McSweeney (2023) | Child (6.92±2.21) | 1/f (FOOOF) | RECORDING, EPOCH<br>3 min length; 2s epoch length;<br>PSD ESTIMATION<br>Welch, 2s Hamming, 50% overlap<br>Single 1-49Hz spectrum per sub<br>REFERENCING<br>mastoid reference (1/2 TD studies)<br>MODEL<br>3-40Hz. peak_width_limits=[1, 8], min_peak_height=0.05, peak_threshold=0.5, max_n_peaks=6 | Global | EOR: 1.53±0.31<br>ECR: 1.77±0.28 | | EOR, ECR | 502(230,272) | Section 3.2, Auth Corr. |
| 13 | Arnett (2022a) …Stein | Child (8.83±1.23) | 1/f (FOOOF) | RECORDING, EPOCH<br>85-120s; continuous segment<br>PSD ESTIMATION<br>Welch, 1s Hamming, 50% overlap<br>REFERENCING<br>online: vertex (Cz); offline: Common average<br>MODEL<br>1-50Hz. peak_width_limits=[2,12], min_peak_height=0.5, peak_threshold=2.0, max_n_peaks=8 | Global | 1.77±0.15 (Median: 1.76) | | EOR | 29(19,10) [b] | Methods, Auth Corr. |
| 14 | Arnett (2022b) … Levin | Child (8.83±1.23) | 1/f (FOOOF) | RECORDING, EPOCH<br>85-120s; continuous segment<br>PSD ESTIMATION<br>Welch, 1s Hamming, 50% overlap<br>REFERENCING<br>Online: vertex (Cz); Offline: Common average<br>MODEL<br>1-50Hz. peak_width_limits=[2,12], min_peak_height=0.5, peak_threshold=2.0, max_n_peaks=8 | Global | 1.77±0.15 (Median: 1.76, range: 0.22-2.30) | | EOR | 29(19,10) [b] | Methods, Auth Corr. |

| | | | | | | | | | | |
|---|---|---|---|---|---|---|---|---|---|---|
| 15 | Peisch & Arnett (2022) | Child<br>(9.40±1.36) | 1/f<br>(FOOOF) | RECORDING, EPOCH<br>85s length; continuous segment<br>PSD ESTIMATION<br>Welch, 1s Hamming, 50% overlap<br>REFERENCING<br>Online: vertex (Cz); Offline: Common average<br>MODEL<br>1-50Hz. peak_width_limits=[2,12], min_peak_height=0.5, peak_threshold=2.0, max_n_peaks=8 | Global<br>Regional | 1.78±0.14<br>Anterior Frontal (AF): 1.79±0.14<br>Frontal (FR): 1.79±0.13<br>Central (CE): 1.75±0.15<br>Parietal (PR): 1.81±0.16<br>Occipital (OC): 1.77±0.22 | | EOR | 29(19,10) [b] | Methods,<br>Auth Corr. |
| 16 | Hill (2022) | Child<br>(9.41±1.95) | 1/f<br>(FOOOF) | RECORDING, EPOCH<br>2 mins ECR, 2 mins EOR; 2s<br>PSD ESTIMATION<br>Hamming window 50% overlap, 2s<br>REFERENCING<br>Online: Cz, Offline: common average<br>MODEL<br>1-40Hz. peak_width_limits=[1,12], min_peak_height=0.0, peak_threshold=2.0, max_n_peaks=8 | Global<br><br>Regional (anterior [A], central [C], posterior [P]) | EOR: 1.65±0.18<br>ECR: 1.81±0.16<br>EOR: A (1.64±0.19), C (1.69±0.19)<br>P (1.68±0.20)<br>ECR: A (1.81±0.17), C (1.85±0.16),<br>P (1.84±0.18) | | EOR, ECR | 139 (72, 67) | Fig 2,<br>Auth Corr. |
| 17 | Trondle (2022) | Child (N=153), Adolescent (N=34), Young Adult (N=3)<br>(10.07±3.39, 5.02-21.67) | 1/f<br>(FOOOF) | RECORDING, EPOCH<br>EO 20s; 1m 40s total, EC 40s; 3m 20s total; 2s epochs; averaged<br>PSD ESTIMATION<br>Welch, 2s sliding windows, 0.25Hz resolution<br>REFERENCING<br>Online: Cz, Offline: common average<br>MODEL<br>peak_width_limits=[0.5,12], min_peak_height=0.0, peak_threshold=2.0, max_n_peaks=inf, aperiodic_mode='fixed' | Regional (Parieto-occipital) | 1.89±0.36 (0.68-2.77)<br>Child: 1.98±0.30<br>Adolescent: 1.58±0.37<br>Young adult: 1.12±0.04 | | ECR | 190<br>(104,86) | Methods,<br>Auth Corr.,<br>Fig 3, App. 4, Supp. 2, 3 |
| 18 | Kwok (2019) | Child<br>4yrs (N=8),<br>5yrs (N=14),<br>6yrs (N=11),<br>(5.60±?.??) | HE | RECORDING, EPOCH<br>3 min EO EC; 60 x 20s epochs<br>REFERENCING<br>Bilateral mastoid average<br>HURST SCALES<br>log(1-19.5s) | Global,<br><br>Channelwise [est] | Median: ~0.09 (EOR)<br>~0.06(ECR)<br>Posterior electrodes: ~0.09 EOR, ECR | | EOR, ECR | 33(?,?) | Fig 6A-C |
| 19 | Smit (2011) | Child<br>(5.27±0.19, 6.79±0.19),<br>Adolescent (16.06±0.55 17.57±0.55),<br>Young Adult (26.18±4.15)<br>(5-50) | HE | RECORDING, EPOCH<br>3-4mins;<br>REFERENCING<br>Earlobes<br>HURST SCALES<br>log windows(1.5-20s) | Channelwise (12 channels) | Child Theta (P3 maxima): 0.77±0.09 (5yrs), 0.76±0.07 (7yrs)<br>Child Alpha (O2 maxima): 0.70±0.09 (5yrs), 0.71±0.08 (7yrs)<br>Child Beta: 0.64±0.09 (5 yrs, Fp2), 0.62±0.08 (7yrs, F8)<br>Adol Theta (Fp1 maxima): 0.72±0.06 (16yrs), 0.72±0.06 (18yrs)<br>Adol Alpha (O1 maxima): | | ECR | 5yrs 366<br>7yrs 378<br>16yrs 426<br>18yrs 387<br>25yrs 396 | Auth Corr.<br>Methods,<br>Fig 3,<br>Table 2 |

| | | | | | | | | | | |
|---|---|---|---|---|---|---|---|---|---|---|
| | | | | | | 0.72±0.10 (16yrs), 0.73±0.12 (18yrs)<br>Adol Beta: 0.64±0.09 (16yrs), 0.66±0.11 (18yrs)<br>YA Theta (F3 maxima): 0.73±0.07 (25yrs)<br>YA Alpha (P4 maxima): 0.75±0.09 (25yrs)<br>YA Beta (O1 maxima): 0.67±0.10 (25yrs) | | | | | |
| 20 | Bruining (2020) | Child (10.30±1.54) | HE | RECORDING, EPOCH<br>3-5 mins ECR;<br>REFERENCING<br>Online: Common mode sense, Offline: common average<br>HURST SCALES<br>2-30s | Global | 0.66±0.04 | | ECR | 29 (14,15)[a] | Supp. Table 1 | Y |
| 21 | McSweeney (2021) | Adolescent (12-17) | 1/f (FOOOF) | RECORDING, EPOCH<br>3 mins ECR, 3 mins EOR; 3s<br>PSD ESTIMATION<br>Hanning taper, resolution 0.33Hz<br>REFERENCING<br>Online: Cz, Offline: Common average<br>MODEL<br>1-45Hz. peak_width_limits=[1,6], min_peak_height=0.0, peak_threshold=2.0, max_n_peaks=4, aperiodic_mode='fixed' | Global (1-45Hz) | $t_1$ (all subjects): EOR (1.21±0.30)<br>$t_1$ (subjects with t1 & t2): EOR (1.21±0.30)<br>$t_1$ (all subjects): ECR (1.33±0.27)<br>$t_1$ (subjects with t1 & t2): ECR (1.21±0.30)<br>$t_2$ (all subjects): EOR (1.10±0.26)<br>$t_2$ (subjects with t1 & t2): EOR (1.11±0.27)<br>$t_2$ (all subjects): ECR (1.16±0.25)<br>$t_2$ (subjects with t1 & t2): ECR (1.17±0.26) | | EOR, ECR | 186 (85,101)<br><br>95 @$t_1$, $t_2$ | Fig 1B, Results | |
| 22 | Ostlund (2021) | Adolescent (13.97±1.28) | 1/f (FOOOF) | RECORD, EPOCH<br>4mins EO, 4 mins EC; unknown<br>PSD ESTIMATION<br>Fourier transform, 0.5Hz increm. 1-50Hz<br>REFERENCING<br>Online: Cz, Offline: average.<br>MODEL<br>2-50Hz. peak_width_limits=[1,8], max_n_peaks=8, aperiodic_mode='fixed' | Global (2-50Hz) | (EOR+ECR/2): 1.80±0.28 (0.92-2.57)<br>EOR: 1.72±0.31 (0.88-2.49)<br>ECR: 1.88±0.28 (0.90-2.65) | | EOR, ECR | 97(53,43) | Table 1 | |
| 23 | Linkenkaer-Hansen (2007) | Adolescent (16.50-19.50) | HE | RECORDING, EPOCH<br>3-6mins; NA<br>REFERENCING<br>Online: Linked earlobes, Offline: Common average<br>HURST SCALES<br>1-20s | Channelwise (Alpha, Beta) | Alpha: (0.70-0.74±0.08-0.11)<br>Beta:(0.61-0.66±0.07-0.09) | | ECR | 390 (196,194) | Table 1 | |

| | | | | | | | | | | | |
|---|---|---|---|---|---|---|---|---|---|---|---|
| 24 | Gao, F. (2017) | Adolescent<br>(18.30±2.80) | HE | RECORDING, EPOCH<br>5 mins; 2 mins<br>REFERENCING<br>Online: Mastoid average<br>HURST SCALES<br>log(1-15s) | Channelwise<br>(Delta-<br>Gamma)$^{est}$ | Alpha: ~0.80<br>Beta: ~0.70 | 0.60<br>0.40 | ECR | 15(15,0) | Fig 2 | |
| 25 | Donoghue<br>(2020) | Young Adult<br>(19.56±1.90) | 1/f<br>(FOOOF) | RECORDING, EPOCH<br>2 min<br>PSD ESTIMATION<br>Welch, 2s windows, 50% overlap<br>REFERENCING<br>Offline: Common average<br>MODEL<br>2-40Hz. peak_width_limits =[1,6],<br>max_n_peaks =6, min_peak_height =0.05,<br>peak_threshold =1.5, aperiodic_mode ='fixed' | Channelwise<br>(Cz) | 1.43±0.25 | | EOR | 16(8,8) | Auth Corr.,<br>Results | |
| 26 | Linkenkaer-<br>Hansen (2001) | Young Adult<br>(20-30) | 1/f (PLE)<br><br>HE | RECORDING, EPOCH<br>20 mins; 120s window<br>PSD ESTIMATION<br>0.005-0.5 Hz<br>REFERENCING<br>Unknown<br>HURST WINDOW<br>Unknown | Global (Alpha<br>[8-13Hz])<br>4-channel avg | PLE ECR: 0.36±0.17<br>PLE EOR: 0.51±0.12<br>HE ECR: 0.68±0.07<br>HE EOR: 0.70±0.04 | | EOR, ECR | 10(9,1) | Results | Y |
| 27 | Muthukumarasw<br>amy & Liley<br>(2018) | Young Adult<br>(23.00±??) | 1/f<br>(IRASA) | RECORDING, EPOCH<br>5 min; 10s segments<br>PSD ESTIMATION<br>Hanning window, FFT length "next $a^2$ after<br>multiplying sub-epoch data length * max(*h*)"<br>REFERENCING<br>Online: Common average, Offline: FCz.<br>MODEL<br>resampling factor *h* 1.1-2.9 (0.05 steps);<br>$\beta_{lf}$ 0.1-2.5Hz<br>$\beta_{hf}$ 20-100Hz | Global,<br><br>Channelwise | $\beta_{lf}$ 1.36(1.12-1.72)<br>$\beta_{hf}$ 1.48(1.18-1.81)<br>$\beta_{lf}$ frontal maxima: 1.72<br>$\beta_{hf}$ central maxima: 1.81 | | ECR | 17(17,0) | Methods,<br>Supp. Fig 7 | |
| 28 | Pathania (2021) | Young Adult<br>(20.88±2.24) | 1/f (PLE)<br>1/f<br>(FOOOF) | RECORDING, EPOCH<br>2 mins; 1s segments<br>PSD ESTIMATION<br>Welch, 50% overlap; FFT 0.977Hz bins,<br>Hamming 50% taper.<br>REFERENCING<br>Online: Left earlobe, Offline: Ear average<br>MODEL<br>2-25Hz. | Global<br>(FOOOF),<br>Regional<br>(FOOOF), | 1.36±0.26<br>F(1.18±0.34), C(1.40±0.28),<br>P(1.46±0.28), O(1.41±0.29) | | EOR | 59(19,40) | Auth Corr. | |
| 29 | Barry (2021) | Young Adult<br>(21.20±3.80) | 1/f PN<br>Slope<br>(PaWNextra) | RECORDING, EPOCH<br>4min EOR; 2min ECR; 2s segments<br>PSD ESTIMATION<br>Hanning window 10% overlap, DFT<br>correction factor 1.0529 | Global<br><br>Channelwise<br>(30 channels) | EOR (session 1, 2 average):<br>1.07±0.33<br>ECR: 1.22±0.38<br>EOR: 0.41-1.50 (Fp1, Cz)<br>ECR: 0.38-1.22 (Fp1, C4) | | EOR, ECR | 20(3,17) | Auth Corr. | |

| | | | | | | | | | | | |
|---|---|---|---|---|---|---|---|---|---|---|---|
| | | | | REFERENCING<br>A1<br>MODEL<br>0.5Hz extrapolated to 2-24Hz,<br>Ln(Power) 1Hz | | | | | | | |
| 30 | Merkin (2023) | Young Adult<br>(22.20±3.90,18–35) | 1/f<br>(FOOOF) | RECORDING, EPOCH<br>2 mins; 2s<br>PSD ESTIMATION<br>Welch, 2s Hamming window, 50% overlap<br>REFERENCING<br>Online: Common average<br>MODEL<br>2–40 Hz. aperiodic_mode='fixed' mode.<br>peak_width_limits = [1,12],<br>maximum_n_peaks = 6, peak_threshold = 2,<br>min_peak_ height = 0 | Global $^{est}$<br>Regional | ~1-2.1<br>~ range 1.3-1.6 YA | | ECR | 85(37,48) | Sections 2.1,<br>3.1, Supp. S5 | |
| 31 | Ke (2022) | Young Adult<br>(22.29±2.28) | 1/f<br>(FOOOF) | RECORDING, EPOCH<br>Unknown; 2mins<br>PSD ESTIMATION<br>FFT 0.5-30Hz.<br>REFERENCING<br>Online: FCz, Offline: Mastoid average<br>MODEL<br>peak_width_limits = [1.0,5.0], max_n_peaks = 6, min_peak_height = 0.1, peak_threshold = 1, aperiodic_mode = 'fixed' | Global,<br>Regional<br>(Frontal,<br>Central,<br>Parietal,<br>Occipital) | Global (1.84±0.34)<br>Frontal (1.99±0.35)<br>Central (1.84±0.34)<br>Parietal (1.76±0.37)<br>Occipital (1.67±0.52) | | EOR | 90(44,46) | Table 1,<br>Auth Corr. | |
| 32 | Smit (2013) | Young Adult<br>(22.40, 21–25) | 1/f (PLE)<br>HE | RECORDING, EPOCH<br>Unknown; 6 mins<br>PSD ESTIMATION<br>Welch, 75% overlap, Hanning envelope 64s.<br>REFERENCING<br>Common average<br>MODEL<br>0.0156-2Hz<br>HURST WINDOW<br>0.5-64s (0.0026-.067Hz) | Channelwise<br>(Alpha [9-13Hz])<br>CP3 | Maxima (both): central midline, scalp ranges Hurst (0.70-0.80), 1/f (0.20-0.40)<br>PLE = 0.43<br>HE = 0.66 (Range: 0.66-1.04) | | EOR | 39(11,28) | Fig 1B/C,<br>Auth Corr. | Y |
| 33 | Zsido (2022) | Young Adult<br>(22.48±3.79) | 1/f<br>(FOOOF) | RECORDING, EPOCH<br>11min; continuous recording<br>PSD ESTIMATION<br>Welch, 4s window, 50% overlap<br>REFERENCING<br>Online: Right mastoid, Offline: Common average<br>MODEL<br>1-40Hz. FOOOF parameters unspecified. | Global $^{est}$ | ~1.40 | | ECR | 31(?,?) | Methods | |
| 34 | Immink (2021) | Young Adult<br>(22.67±3.85) | 1/f<br>(IRASA) | RECORDING, EPOCH<br>5 mins ECR, 5 mins EOR; 30s ECR<br>PSD ESTIMATION<br>IRASA | Global | 2.06±0.13 (range: 1.82-2.48) | | ECR | 45(22,23) | Section 3.1,<br>Auth. Corr. | |

| | | | | | | | | | | |
|---|---|---|---|---|---|---|---|---|---|---|
| | | | | REFERENCING<br>Online: FCz, Offline: Mastoid average<br>MODEL<br>0.1-40Hz, resampling factor 1.1-1.9 (0.05 steps) | | | | | | |
| 35 | Pathania (2022) | Young Adult (23.29±3.47) | 1/f (FOOOF) | RECORDING, EPOCH<br>2 mins; 1s segments<br>PSD ESTIMATION<br>Welch, 50% overlap; FFT 0.977Hz bins, Hamming 50% taper.<br>REFERENCING<br>Online: right ear.<br>MODEL<br>2-25Hz. peak_width_limits=[1.0, 8.0], max_n_peaks=8, min_peak_height=0.05, peak_threshold=2.0, aperiodic_mode='fixed' | Global (2-25Hz), Regional | 1.17±0.23<br>F(1.20±0.25), C(1.22±0.27), P(1.09±0.28), O(0.96±0.28) | | ECR | 21(11,10) | Section 4.1, Figure 2B, Auth Corr. |
| 36 | Cross (2022) | Young Adult (25.00±7.13) | 1/f (FOOOF, IRASA) | RECORDING, EPOCH<br>2 mins ECR, EOR; 13.2s segments<br>PSD ESTIMATION<br>(FOOOF-only) Welch, Hann-tapered, zero-padded to 2048 points<br>REFERENCING<br>Online: FCz, Offline: mastoid average<br>MODEL<br>*IRASA*<br>$h$ = 1.1-1.95 (0.05 steps)<br>*FOOOF*<br>1-35Hz. peak_width_limits=[1, 12], maximum_n_peaks=infinite, min_peak_height=0, peak_threshold=2.0, aperiodic_mode= 'fixed' | Global | *IRASA* ECR: 1.11±0.30<br>*IRASA* EOR: 1.08±0.31 | | EOR, ECR | 35 (18,17) | Auth Corr. |
| 37 | Nakao (2019) | Young Adult (19.57±?? 18-21) | HE | RECORDING, EPOCH<br>5 mins; 30s segments<br>PSD ESTIMATION<br>75% overlap<br>REFERENCING<br>Offline: Common average<br>HURST WINDOW<br>log(1-30s) | Alpha [8-13Hz] | FCz: 0.75±0.12<br>Min (T7, 0.74±0.12)<br>Max (O1, 0.80±0.13) | 0.50 (0.48-0.60) | ECR | 23(11,12) | Fig 5, Section 3.3, Auth Corr. |
| 38 | Natarajan (2004) | Young Adult (20.00±3.00) | HE | RECORDING, EPOCH<br>20 mins; 5 mins<br>REFERENCING<br>Unknown<br>HURST SCALES<br>Unknown | Global (1-50Hz) | 0.29±0.06 | -0.42 | ECR | 30(15,15) | Table 1 |
| 39 | Liu, S (2022) | Young Adult (20-30) | HE | RECORDING, EPOCH<br>4 mins ECR, 4 mins EOR; 5s segments<br>PSD ESTIMATION | Channelwise [est] (Broadband) [0.5-120Hz] | EOR<br>~0.80-0.82 | EOR<br>0.60-0.64 | ECR, EOR | 26(?,?) | Fig 5 |

| | | | | | | | | | | | |
|---|---|---|---|---|---|---|---|---|---|---|---|
| | | | | Welch, Hamming window, 50% overlap<br>REFERENCING<br>Online: Linked left mastoid (M1), Offline: Average mastoid<br>HURST WINDOW<br>Unknown | | | | | | | |
| 40 | Sleimen-Malkoun (2015) | Young Adult (22.70±1.60, 18.80-25.10) | HE | RECORDING, EPOCH<br>1.5 min ECR, EOR; 10s segments<br>PSD estimation<br>2500pt Hanning window; 4096pt (zero-padded)<br>REFERENCING<br>Online: left mastoid, Offline: average mastoid<br>HURST WINDOW<br>4-50ms, 16-200ms, 4ms steps | Global (0.5-100Hz) | 1.69<br>Higher for posterior vs midline | 2.38 | ECR | 31(17,14) | Fig 4 | Y |
| 41 | Irrmischer (2018) | Young Adult (25.00±6.20) | HE | RECORDING, EPOCH<br>5mins ECR, 5mins EOR<br>REFERENCING<br>Online: Cz, Online: Common average<br>HURST WINDOW<br>5-30s (delta, theta)<br>2-30s (alpha)<br>1-30s(beta)<br>1-30s (gamma) | Global (Delta [1-4Hz], Theta [4-8Hz], Alpha [8-13Hz], Beta [13-45Hz]) | ECR (N = 57)<br>Theta: 0.66±0.01<br>Alpha: 0.71±0.01<br>Beta: 0.66 ± 0.01<br>EOR (N = 23)<br>Theta: 0.69±0.02<br>Alpha: 0.75±0.02<br>Beta: 0.70 ± 0.01 | ECR<br>0.32<br>0.42<br>0.32<br>EOR<br>0.38<br>0.50<br>0.40 | EOR, ECR | 57(22,35) | Results | |
| 42 | Bornas (2013) | Young Adult (24.61±7.03) | HE | RECORDING, EPOCH<br>8 mins; 1 min segments (EOR, ECR)<br>REFERENCING<br>Linked earlobes<br>HURST WINDOW<br>0.1-0.6s (broadband),<br>1-6s (narrowband) | Regional (theta [3-7Hz], alpha [8-13Hz], broadband [1-40Hz]): Central [C], Parietal [P], Occipital [O]) | Theta<br>C(0.75±0.07), P(0.76±0.07), O(0.74±0.07)<br>Alpha<br>C(0.76±0.07), P(0.80±0.08), O(0.85±0.10)<br>Broadband<br>C(0.85±0.07), P(0.86±0.06), O(0.88±0.06) | Theta<br>0.50,0.52<br>0.48<br>Alpha<br>0.52, 0.60<br>0.70<br>Broadband<br>0.70, 0.72<br>0.76 | EOR, ECR average | 56(20,36) | Table 1 | |

**A**

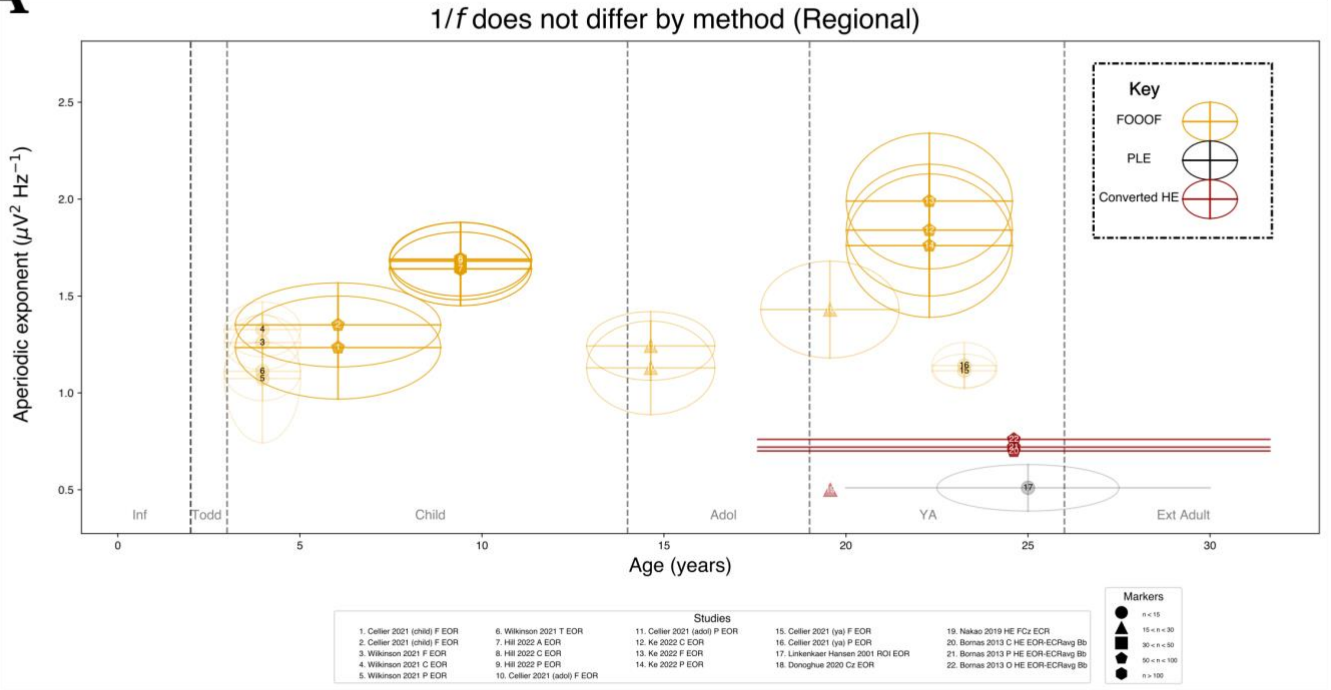

**B**

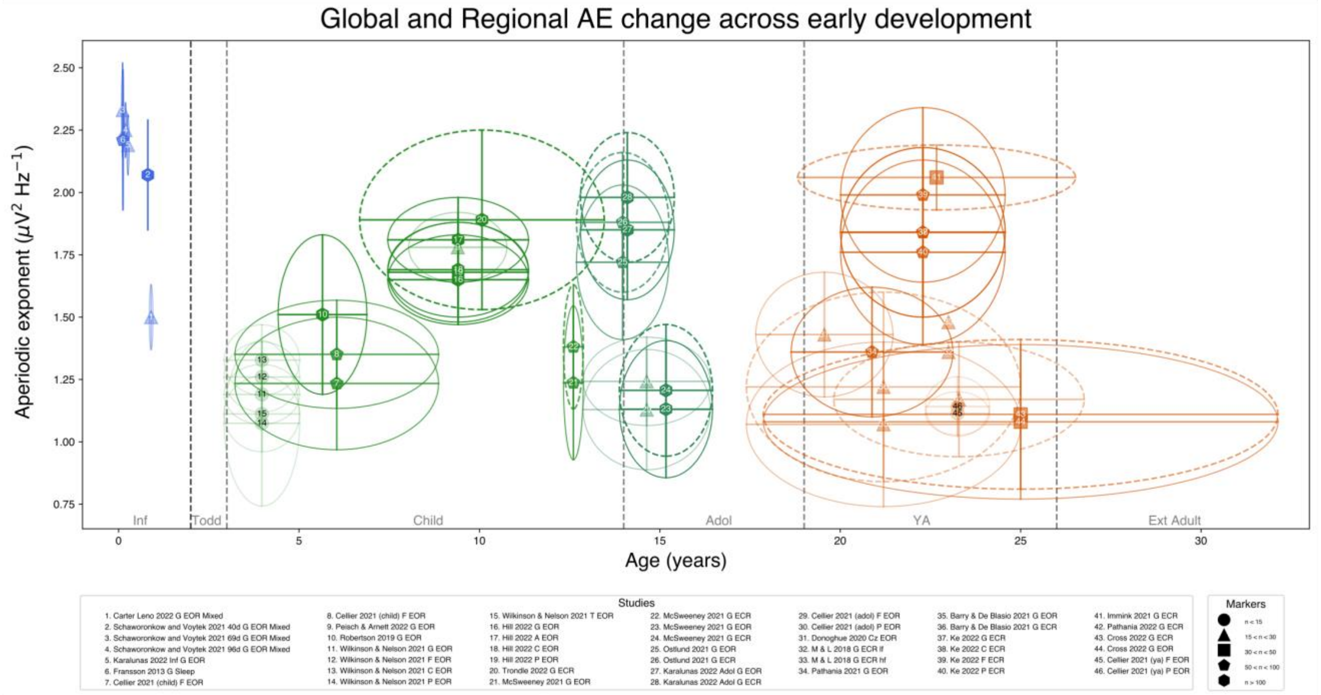

***Supplement V.*** *Regional AE do not differ by method.*

(A) ROI AE data is sparse, yet towards adulthood some converted HE are within adequate scope of overlap, whilst others, similar to PLE exhibit are more broadly dispersed. (B) Studies are colour-coded by lifespan stage, and results for both EOR and ECR are shown for all available studies. Age related changes are apparent during infancy, but not across prolonged development.
